# The virus integrations in gut microbes associated with dysbiosis of microbial community in tumorigenesis of colorectal carcinoma

**DOI:** 10.1101/867291

**Authors:** Boyu Cui, Xiaodan Wang, Wenqiu Xu, Caihong Zheng, Jun Cai

## Abstract

Patients with colorectal cancer (CRC) have a different gut microbial and viral communities from healthy individuals. But little is known about the ways and functions of interaction of virus-bacteria, let alone its correlation with the aetiology of CRC. In this study we aimed to identify the association between the genetic integration of virus-bacteria and the expansion of some microbial population during tumorigenesis of human colorectum. Using a gut metagenomics sequencing data of healthy controls, advanced adenoma and carcinoma patients, to our knowledge, we demonstrate for the first time that the viral genetic integrations in gut microbes tend to occur in CRC patients and are potentially associated with the carcinogenesis. We found that almost all of the genetic integrations were happened between bacteriophages and bacteria, which could be influenced by the abundance of the phage communities. Importantly, the integrations of phage-carried positive effective genes offered selective advantages to the commensal and potential pathogenic bacteria, as a result, potentially led to a microbial dysbiosis along with the increasing bacterial diversity in carcinoma patients. Consequently, our work opens a new way to understand the carcinogenicity in complex intestinal ecosystems.

## Introduction

Colorectal cancer (CRC) is among the top three most common cause of cancer mortality in the world. Most of the CRC cases are referred to as the order of adenoma-carcinoma that could progress into malignant forms [1–3]. The accumulation of genetic mutations for many years is the most important reason for the development of CRC, which is affected by many factors such as lifestyle and environment including the gut microbiota [4–6]. The human gut microbiome is composed of a large number of bacterial cells, along with a minority of virus, archaeal and eukaryotic cells, together forming a very complex intestinal microbial ecosystem [7]. The virus is primarily made up of bacteriophages and also includes rarer eukaryotic viruses and endogenous retroviruses [8–10].

Accumulating data have demonstrated that the composition of gut microbiota in CRC patients are distinct from healthy individuals [11, 12]. On the one hand, emerging evidences have indicated that several bacteria, such as *Bacteroides fragilis* [13], *Fusobacterium nucleatum* (*F. nucleatum*) [6, 14] and a strain of *Escherichia coli* [4], are more abundant in the gut of CRC patients or CRC tissues and have been found to be associated with CRC development by promoting colorectal carcinogenesis. These microbes can accelerate carcinoma development through secretion of oncogenic virulence factors [15], generation of carcinogenic microbial metabolites [16, 17] or recruitment of immune cells [18, 19] and so on. On the other hand, an increasing number of studies suggest that the human intestinal microbiota contributes to CRC via the influence of the whole microbial community, especially the dysbiosis of the bacterial population that is a result of symbiotic, commensal, and pathobiont interactions among microorganisms [20, 21]. Using large-scale metagenomic sequencing, recent studies have demonstrated that microbial dysbiosis plays an important role in disorder of immune system and has been shown to modulate the pathological functions associated with CRC, including cell proliferation, apoptosis and immune response [17, 22–24].

However, given the close connection between microbiome and human health, a better understanding of what factors drive bacterial diversification and dysbiosis in the gut is required. Although several studies have considered the effects of different factors such as diet, age, geography and host genetics on structural variation of the gut microbiome [25, 26], the effects of virus-bacteria interaction in the gut have been paid more and more attention by researchers [27]. Recently, the metagenome-wide association study of gut virome in CRC has demonstrated that virome profile differentiated individuals at healthy controls, early and late stages of CRC patients [28]. Moreover, statistical comparison between gut phage and bacterial communities provide the evidences that there could be a close and dynamic relationship between these phages and bacterial population. Their significant correlation suggests that phages, to a certain extents, play roles in reshaping microbial ecosystems and consequently affect the health or disease status of the individuals. Undoubtedly, the form and function of virus-bacteria interaction in the gut are potentially manifold. As phages have the ability to help to facilitate the release of bacteriocin [29], improve the metabolism ability [30], and enhance the biofilm formation [31], they can increase the adaptive and competitive ability of bacteria within the gut ecosystem. Importantly, bacteriophages are able to change microbial population by predating on them resulting in host lysis or by accelerating their adaptation through horizontal gene transfer [32, 33]. Nevertheless, the effects of virus-bacteria interaction at this level in the gut have not been received much research attention. Consequently, we have little knowledge of how potentially important interaction of bacteria-bacteriophage could impact on the communities and consequent functionality of the gut microbiota.

We therefore performed a metagenomic study of gut microbiota in healthy controls, advanced adenoma and carcinoma patients with a particular focus on the interaction of virus-bacteria. Using a strict pipeline of bioinformatics, we found, for the first time, that the viral genetic integrations in the microbial community were associated with the colorectal tumorigenesis. Firstly, we identified that the integrations between virus and bacteria communities, actually mainly dominated by phage and bacteria, were generally existent phenomenon in human gut, especially, in colorectal carcinoma patients. And these integrations could be associated with the abundance of the phage communities. Secondly, the integrations of phage-carried positive effective genes offered selective advantages to the commensal and potential pathogenic bacteria, which potentially lead to a microbial dysbiosis along with an increasing bacterial diversity in carcinoma patients. Thirdly, the network of high frequent phage-bacteria integrations potentially played a key role in the development of phage therapy as an alternative to antibiotics. These findings open a new research opportunity for understanding the microbiome associated pathogenesis of colorectal carcinoma.

## Materials and methods

### Study cohorts information

The metagenomic shotgun sequencing data for all 112 samples with high-quality sequencing reads (we chose 5 GB per sample on average) have been downloaded from the EBI database under the accession code ERP008729. The age of data from Caucasians were between 43-86 years in the initial analysis, including 34 healthy controls (12 females and 22 males), 34 advanced adenoma (17 females and 17 males) and 44 carcinoma (17 females and 27 males) patients as shown in supplemental table (Table S1) [4].

### Data preprocessing

Sequences were quality filtered, decontamination and demultiplexed (Fig. S1). In detail, the adapters and low-quality bases (PHRED q < 25) of metagenomic reads were trimmed using Trimmomatic (v.0.36) [34]. The reads were cut each sequence with a 4-base sliding window trimmer that required minimum average quality scores of 15 and removed any sequence that had 36 bases or fewer. We clipped any fragments of adapter sequences in paired-end reads and then decontaminated the post-quality-trimmed metagenomic reads by performing BWA (v.0.7.15) [35] alignment with the indexed mouse (mm10) and human (hg38) reference genomes with default settings, which were considered potential sources of human habitat- or sample-associated contaminants for taxonomic classification of microbial metagenomic sequences. Afterwords, sequences were demultiplexed using Picard MarkDuplicates (v.2.2.1) (http://broadinstitute.github.io/picard/) by identifying the start coordinates and orientations of both reads of a read pair. Within a set of duplicate read pairs, the read pairs with the highest base qualities were retained with the others marked as PCR duplicates. These final reads constituted the cleaned metagenomic reads sets and were analyzed in next steps.

### Identifying the potential genetic integration

Using BWA-aln (v.0.7.15) with default parameters, we identified paired-end reads with one read aligning exclusively to all complete bacterial genomes downloaded from the NCBI RefSeq database (accessed on March 7, 2018) and the integrated gene catalogue of the human gut microbiome (IGC) database, which was a high-quality reference catalog of 9,879,896 gut microbial genes [36], while the other read aligning to the complete virus genomes downloaded from NCBI taxonomic database. The statistical putative virus reads inserted into the bacterial genomes all were supported by at least two pair-end reads. Genus and species taxonomic assignments were parsed from the output using in-house perl scripts. The taxonomic ranking of genus or species were identified according to the NCBI taxonomy database and the two or more sequencing reads supported one integration event should be the same. To reduce possible falsepositive results, further filtering steps were performed.

### Filtering the false-positive

For the filtering of the potential false-positive reads, BLASTn and BLASTp were used to validate each read of the pair as specific for bacteria or virus using the NT and NR database (accessed on March 14, 2018). Both potential bacterial and viral reads were searched against NT and NR database using BLAST (v.2.6.0+) [37] with the default parameters and an e-value cutoff of 10^−5^. Reads identified as bacteria in the initial BWA screen were required to match bacteria in NT and NR and not had a best match to virus and other non-bacterial species. Similarly, the same process was repeated on reads identified as virus. The taxonomic assignment for sequencing reads was determined by the taxonomy ID of the top BLAST result. After the filtering of reads, we calculated the last common ancestor (LCA) of all hits based on the congruent taxonomy for all genomes with mappings as described in previous study [38] and counted the amounts of integration in each sample of the three cohorts.

### Taxonomic classification and abundance estimation

The microbial composition in each sample was identified using Kraken software (v.2.0.8-beta) [39]. Briefly, we assigned the cleaned metagenomic reads to microbial taxa using the k-mer based algorithms as implemented in the Kraken taxonomic annotation pipeline. The search database used the MiniKraken DB_8GB database that was constructed from complete bacterial, archaeal, and viral genomes in RefSeq. Each k-mer in a sequencing read was mapped to the lowest common ancestor of all reference microbial genomes that had exact k-mer matches. Reads mapping to virus or bacteria were counted to calculate relative abundances of certain group in each sample. Metagenomic abundance tables were normalized by the dividing individual taxon sequence counts by the total assigned sequence count in a sample using the vegan R package [40].

### The α-diversity analysis of microbiome

The α-diversity (Shannon diversity) index within-samples was calculated using diversity and vegdist functions of Vegan R package (v.2.5.6) in R (v.3.6.1) for abundance matrices generated. Briefly, the taxonomic matrices of bacteria at genus level and virus at species level were used as inputs, where each element was the number of virus or bacteria from a given sample. The variation of dissimilarities were compared among plaques of healthy controls, advanced adenoma and carcinoma patients. Additionally, *p values* were calculated using a Mann-Whitney U test.

### Functional annotation

The reads supported integration events were functionally annotated using eggNOGmapper (v.4.5.1) [41] and the eggNOG database with default parameters using the specific datasets. Statistics profiles at eggNOG, KEGG orthologous group were created. Using the in-house perl scripts and the KEGG database that included bacterial and viral genes annotated in reference genomes, we could infer the functions of bacterial and viral fragments respectively and counted the amounts of genes of each KEGG terms in all the samples of three cohorts.

### Data visualization

The network was displayed using Cytoscape (v.4.5.1) [42] and related algorithms, which showed a one-to-one genetic integrated correspondence from phages to bacterium. The genetic integration datasets were imported from Excel spreadsheets via Cytoscape Table Import functionality and displayed when clicking on a node (biomolecule) or on an edge (link) of the network. Heatmap figures were plotted using pheatmap package (v.0.7.7) using dist and hclust functions in R (v.3.6.1). The bubble charts and histograms were generated in the R software, too.

### Statistical analysis

Statistical modeling and correlation analysis were performed using the GraphPad Prism (v8.0.2) and R (v.3.6.1) software.

## Result

### The genetic integration as a way of virus-bacteria interaction tended to occur in carcinoma

To investigate the interaction between the virus and bacteria in the gut of different disease status, we downloaded the metagenomic shotgun sequencing in cohort of 112 faecal samples with high-quality sequencing reads including 34 healthy controls, 34 advanced adenoma, and 44 carcinoma patients, respectively (Table S1) [4]. Sequences were demultiplexed, quality filtered, and decontamination (Fig. S1). Then we identified the genetic integrated paired-end reads and calculated the last common ancestor (LCA) of all hits as described in the materials and methods. In order to assure the reliability, the statistical putative viral reads inserted into the bacterial genomes all had at least 2x sequencing coverage and filtered the potential false positive reads by blast. After the processing of reads, we counted the amounts of genetic integration in each sample of the three cohorts (Fig. S1). The result showed that the frequency of the integrations was higher in advanced adenoma and carcinoma patients than healthy controls (* P = 0.01−0.05, ** P < 0.01, Wilcoxon rank-sum test, Fig. 1a). To further describe the interaction, we identified the majority of bacterial genera related to the potential integration. A number of genera *Escherichia*, *Enterobacter* along with *Enterococcus*, *Shigella* and *Bacteroides* had the highest frequency of genetic integration in the cohort, most of whom more likely tended to occur in carcinoma samples compared with healthy and advanced adenoma samples (Fig. 1b). More importantly, several species in the genera including *Bacteroides, Escherichia*, *Streptococcu*s and *Enterococcus* could promote colorectal carcinogenesis through direct interaction with host cancer cells [43], secretion of oncogenic virulence factors [44], generation of carcinogenic microbial metabolites [16] and reshaping immune system [45, 46].

**Fig. 1.**
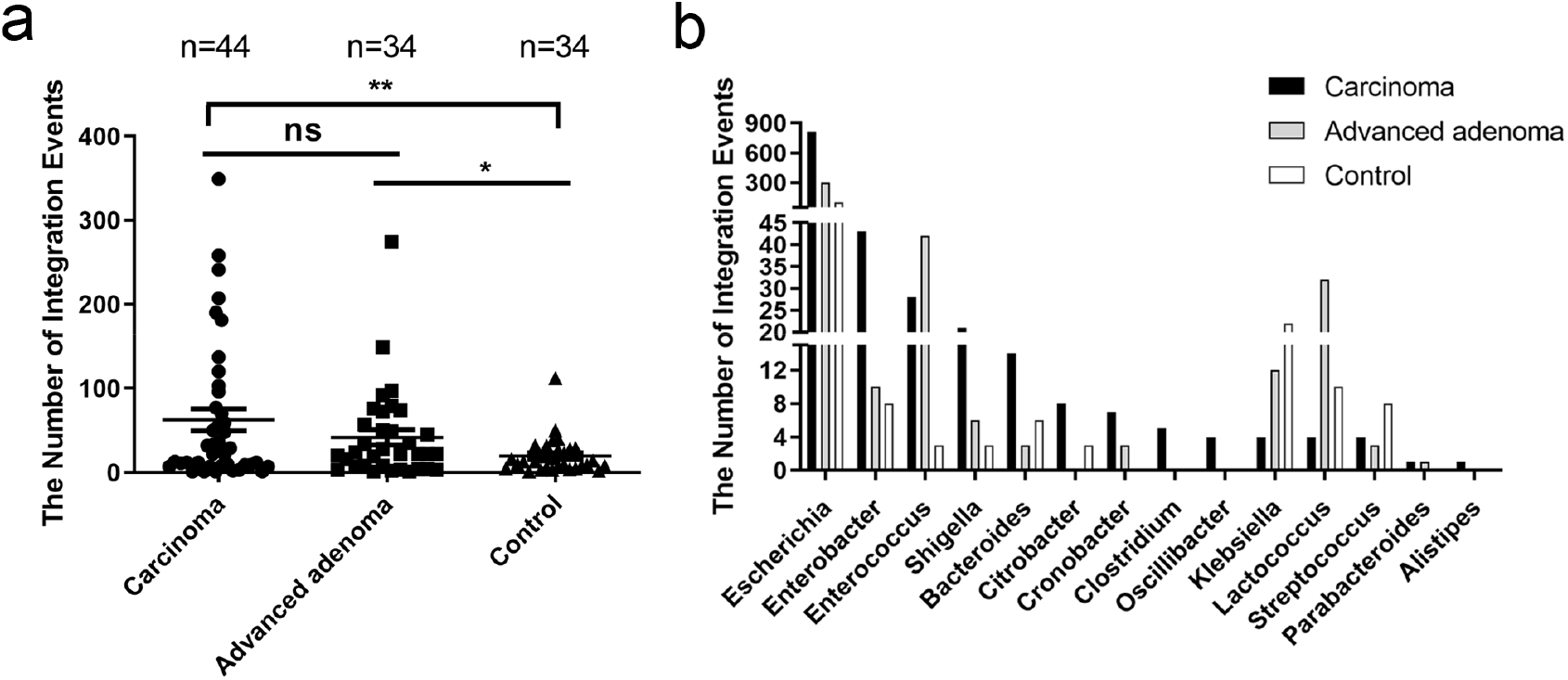
The number of viral genetic integrations into bacterial communities in healthy controls, colorectal carcinoma and advanced adenoma patients. (a). The significant difference of integration events from viruse to bacterium in the three groups are shown. Statistical significance was assessed by two-sided Wilcoxon ranksum test; * P = 0.01-0.05, ** P < 0.01. (b). The frequency of bacterial genera involved in the genetic integration. The bar-plot depicts the frequency of integrations are different at bacterial genus level in the three groups.

Based on the genetic integrations identified in this study, we constructed a virus-bacteria genetic interaction network in the human gut of three groups, respectively (Fig. S2a, b and c). As shown, almost all of the integrations happened between bacteria and bacteriophages that were the main component of virome in human gut [47]. In total, we identified 1490, 428 and 270 phage-bacteria integrations between 283 viral species and 24 bacterial genus, corresponding to 662, 333 and 113 unique integrations in carcinoma, advanced adenoma patients and healthy controls, respectively. Obviously, the transkingdom interactions are more prevalent in carcinoma and advanced adenoma patients than in healthy controls. Besides, we found most of the phage-bacteria integrations were host-specific, however, a small fraction of phages were connected with two or more genera of bacteria, which indicated that they should follow a one-to-many pattern (Fig. S2). Consequently, these results suggest that the genetic integration from virus to bacteria is a generally existent phenomenon in the human gut, especially in carcinoma patients, which potentially play a regulatory role for the bacterium including same pathogens.

### The relative abundance of the phage community associated with the frequency of virus-bacteria integration

In order to determine whether the relative abundance of the bacteria and virus affect the genetic integration or not, the constitution of bacterial and viral communities from the healthy controls, advanced adenoma and carcinoma patients were analyzed. The overall microbial compositions for each groups at the genus level were shown (Fig. S3a). We found that the microbial relative abundance profiles of gut microbiome were different in the three cohorts, and the genera of *Bacteroides*, *Faecalibacterium*, *Blautia*, *Bifidobacterium* as well as *Anaerostipes* had higher relative abundance in the three groups. However, the highest frequency of genetic integrated genera were *Escherichia*, *Enterobacter* along with *Enterococcus*, *Shigella* and *Bacteroides* (Fig. 1b and Fig. S2). These findings show that the relative abundance of the bacteria isn’t the decisive factor for the genetic integration. Nonetheless, as expected, almost all of the viral genetic integrations into their hosts were occurred in three families including *Siphoviridae, Podoviridae and Myoviridae* (Table S2, Fig. S2, Fig S3b, c and d), all of whom were the most abundant viral families in all cohorts consistent with previous report (Fig. 2a) [47]. In addition, we observed the positive correlations between the relative abundance of the three viral families and the frequency of genetic integration with their hosts (Fig. 2b, c and d, Spearman correlation coefficient, r = 0.227, 0.362, 0.410, respectively, P < 0.05). As a result, we demonstrate that the relative abundance of viral communities as an important factor influence the frequency of the virul genetic integrations in their microbial hosts.

**Fig. 2.**
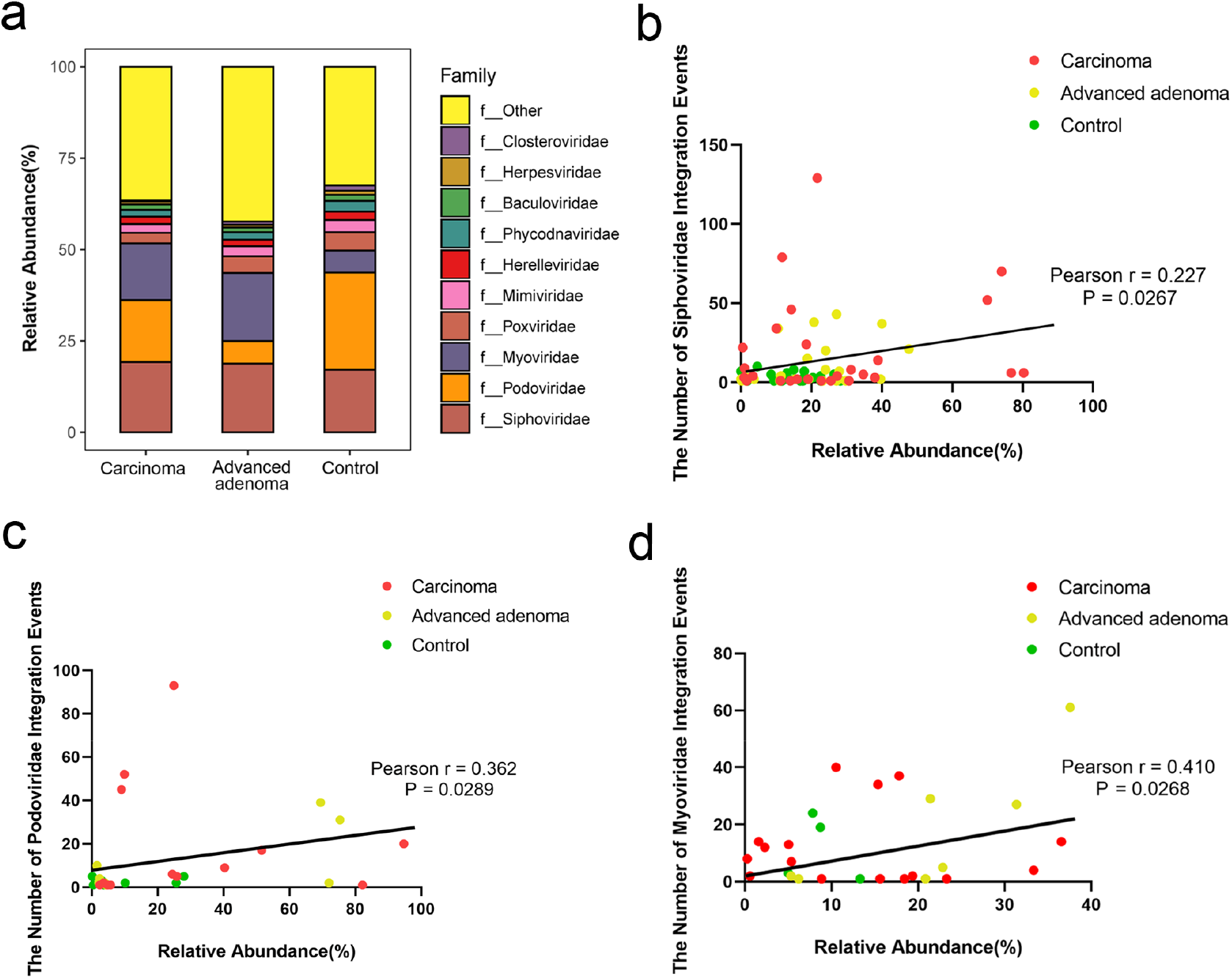
The correlation between the relative abundance of dominant viral families and the frequency of their integrations. (a). The relative abundance of viral families in the three groups including carcinoma, advanced adenoma patients and healthy individuals. Different structures of gut virus at family level are shown. Three families including *Siphoviridae*, *Podoviridae* and *Myoviridae* were found to be significantly higher abundance and almost all of the integrated viral species were occurred among them. (b) Correlation between the abundance of *Siphoviridae* and frequency of its integrations. Each dot represents one sample (n = 73). (c). Correlation between the abundance of *Podoviridae* and frequency of its integrations (n = 29). (d). Correlation between the abundance of *Myoviridae* and frequency of its integrations (n = 28). Line indicates linear regression. Spearman correlation coefficients are shown.

### The integration of phage fragments increasing the successful subpopulation of bacteria

We next asked whether the genetic integrations from phage to bacteria could affect the structure of bacteria in different disease status. First of all, the trans-kingdom integration between bacteriophages and their bacterial hosts were calculated. Heatmap showed the five highest frequency of gut bacterial genera with genetic integration, including *Escherichia*, *Enterobacter*, *Enterococcus*, *Shigella* and *Bacteroides*, in the healthy controls (Fig. 3a), advanced adenoma (Fig. 3b) and carcinoma patients (Fig. 3c). Then to describe the detail effects of integration, the correlation between the integration frequency of five bacterial genera and their relative abundance were identified in each individuals of the three groups. As showed, the integration frequency of *Escherichia* was positively correlated with its relative abundance in carcinoma (Spearman r = 0.64, *P value* < 0.001, Fig. 3d) and advanced adenoma patients (Spearman r = 0.516, *P value* < 0.05), but not in healthy control (Spearman r = 0.075, *P value* > 0.05). As expected, the integration frequency of the other four bacterial genera, which included *Enterobacter*, *Enterococcus*, *Shigella* and *Bacteroides*, were also positively correlated with their abundance in carcinoma patients, however, there were no changes or negative correlations with the relative abundance in healthy control (Fig. S4a, b, c and d). These results show that the genetic integrations display a trend towards increasing the subpopulation of bacteria inserted with phage genetic fragments, especially in carcinoma samples. According to these findings, we speculate that it could contribute to a close relationship between the population diversity of bacteria and virus. To test this idea, the α-diversity (Shannon index) of virus and bacteria in the three groups were analyzed. A significant difference of the viral a-diversity was observed at the species level among the three groups (*P value* = 0.0146, Kruskal-Wallis test, Fig. 3e), while the bacterial a-diversity at the genus level was not, which was consistent with previous studies (Fig. S5) [4]. As hoped, we found a positive association between the microbial and viral diversity in carcinoma patients (Spearman correlation coefficient, r = 0.411, *P value* = 0.002), however, fewer positive or negative correlations were observed in healthy controls (Spearman correlation coefficient, r = 0.450, *P value* = 0.003) and advanced adenoma patients (Spearman correlation coefficient, r = −0.035, *P valu*e > 0.05, Fig. 3f). Consequently, we suggest that a large and diverse pool of phages genetic fragments can be inserted into the genomes of bacteria and confer a selective advantage to individual bacterial cells in carcinoma patients, resulting in successful subpopulations.

**Fig. 3.**
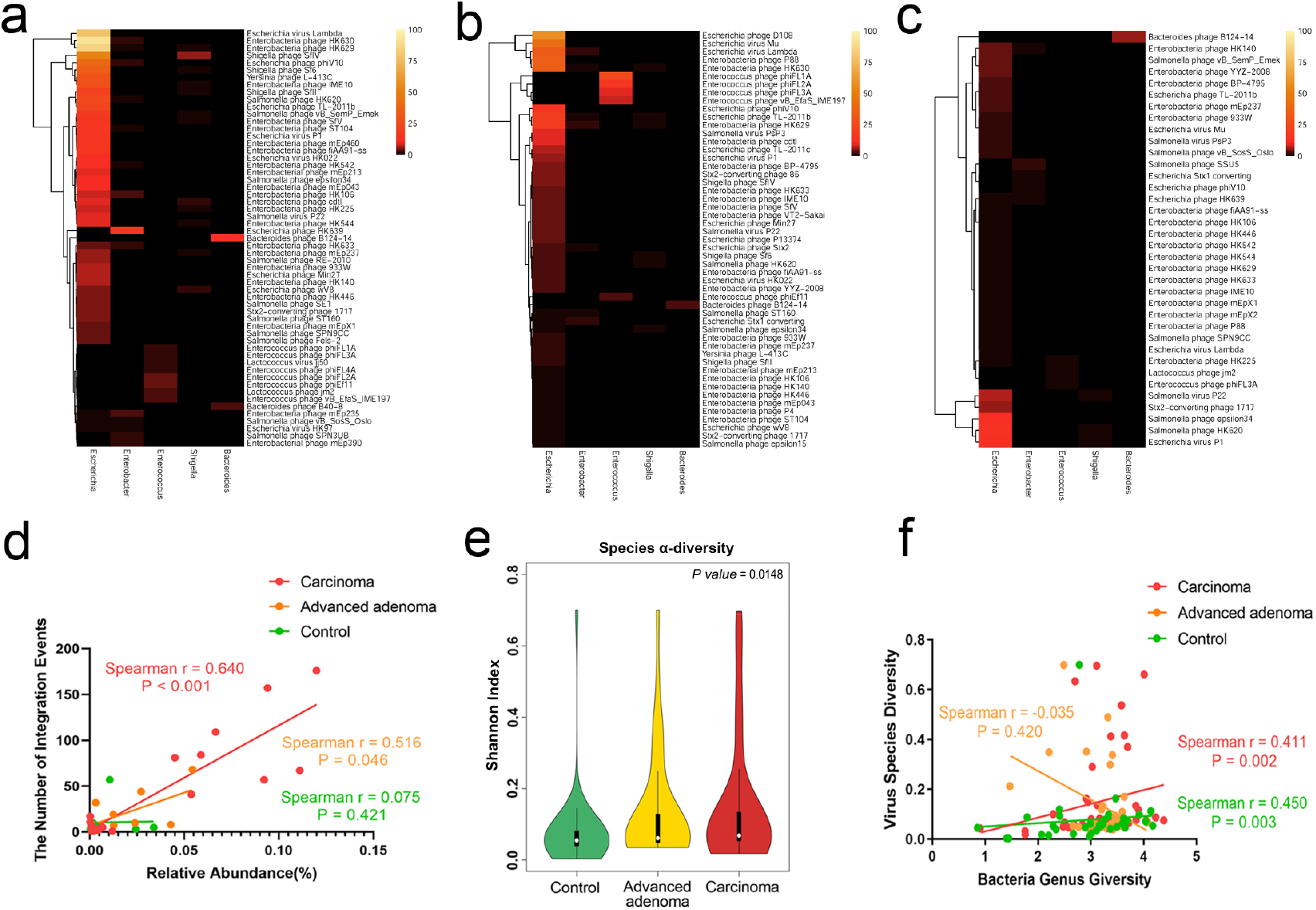
The detail genetic integrations between bacteriophage and the five gut bacterial genera in the different groups. Heatmap figures show the frequency of genetic integrations from bacteriophages to five dominant bacterial genera in the three groups including carcinoma (a), advanced adenoma patients (b) and healthy individuals (c). (d). The relationship between the integrated frequency of *Escherichia* and its relative abundance. The viral genetic integrations of *Escherichia* were positively correlated with its abundance in carcinoma and advanced adenoma patients, while not in healthy control. Each dot represents one sample (n = 32), and the Spearman correlation coefficients are indicated. (e). The viral a-diversity (Shannon index) of the different groups at the species level and *P values* from Kruskal-Wallis tests are shown. (f). Group-specific Spearman rank-order correlation plot showing the relationship between viral and bacterial diversities. Positive correlation was observed between the diversity of bacteria and virus in carcinoma patients; however, negative or no correlations were observed in healthy controls and advanced adenoma patients despite the a-diversities of bacteria wasn’t significant in the three groups. Line indicates linear regression and Spearman correlation coefficients are shown. Healthy controls n = 34, advanced adenoma n=34 and carcinoma patients n=44.

### The integrated phage fragments carrying bacteria-beneficial genes in carcinoma samples

To assess the functional content of the inserted phage fragments, we annotated the shortinsert-size raw reads using the eggnog-mapper (v.4.5.1) [48]. KEGG bubble graphs were plotted against the −log10 (p-value) for demonstrating the functional classification of the predicted integrated phage fragments in the groups of healthy controls, advanced adenoma or carcinoma samples, respectively (Fig. 4a, b and c, respectively). These results displayed that the composite genetic fragments tended to be found in Peptidoglycan biosynthesis, Degradation proteins and Enzymes with EC numbers pathways in all of the three groups. Importantly, we also observed the pathway terms specifically enriched in Replication and repair, DNA replication, Cell cycle and Defense system in carcinoma patients, while not found in advanced adenoma patients and healthy controls. These inserted viral fragments possibly enable to increase metabolism, competitiveness and cell activity (such as proliferation, differentiation and survival) in diverse ways in the host bacteria of carcinoma patients. On the one hand, direct benefits of increased metabolism and cell activity confer the selective advantages to individual cells, resulting in successful subpopulations. On the other hand, these phage-encoded inserted fragments possibly enable hosts to become pathogenic and promote the host cells to form and accumulate biofilm by attachment to neighboring cells, which is considered beneficial to the local bacterial population.

**Fig. 4.**
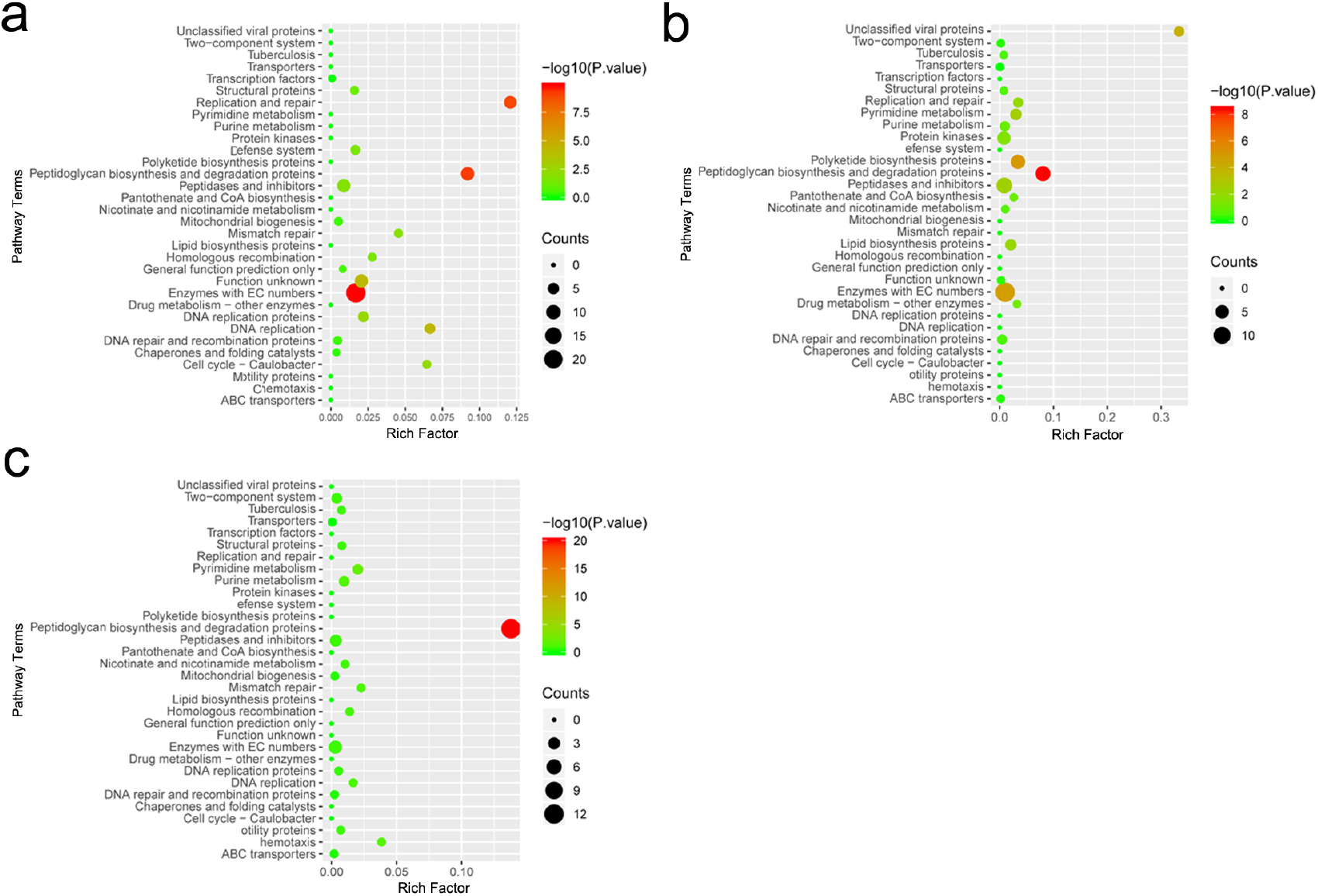
The analysis of Kyoto Encyclopedia of Genes and Genome (KEGG) pathway terms. KEGG bubble charts plotted against the −log10 (p-value) for demonstrating the functional classification of the predicted viral fragments inserted into bacterial chromosomes in healthy controls, colorectal adenoma or carcinoma samples, respectively. Size of bubbles represents the number of proteins enriched for each of KEGG pathway terms; The dark of bubble color represents the *p value* and the X-axis label represents the rich factor (rich factor = amount of inserted genes fragments enriched in the pathway/amount of all genes in background gene set in a certain signaling pathway). The results showed the KEGG pathway terms mainly enriched in Replication and repair, DNA replication, Cell cycle and Defense system in carcinama patients while not in advanced adenoma patients and healthy controls.

### The microbial dysbiosis along with an increasing diversity appearing in colorectal carcinoma patients

In order to analyze the effect of the genetic integrations on the whole microbial community, we calculated the microbial dysbiosis index (MDI), which was defined as calculating the log of [total abundance in organisms increased in carcinoma patients] over [total abundance of organisms decreased in carcinoma patients] for all samples [49]. In brief, we combined the five (the taxon within the genera *Fusobacterium*, *Escherichia*, *Bacteroides*, *Alistipes* and *Porphyromonas* enriched in CRC [13, 50–53]) and three (the taxon within the genera *Bifidobacterium*, *Roseburia* and *Blautia* decreased in CRC [6, 54, 55]) genera that were generally reported and enriched in the carcinoma patients and healthy controls, respectively. As a result, the gut microbiome of carcinoma patients had a higher MDI than that of individuals including advanced adenoma patients and healthy controls (*P value* < 0.0001, Kruskal-Wallis test, Fig 5a). Besides, the MDI showed a positive correlation with the α-diversity of bacteria in carcinoma (Spearman r = 0.292, *P value* = 0.027), while not in advanced adenoma patients (Spearman r = −0.160, *P value* = 0.182) and healthy controls (Spearman r = 0.134, *P value* = 0.224, Fig 5b). In short, these results demonstrate that the carcinoma patients have a state of dysbiosis of the gut microbiome along with an increasing bacterial α-diversity. In addition, a number of genera, including *Escherichia*, *Enterococcus*, *Bacteroides*, *Clostridium* and *Alistipes*, had higher frequency of viral genetic integration in carcinoma patients compared with healthy controls and advanced adenoma individuals (Fig. 1b). Importantly, some species of bacteria in the genera are often reported enriched in CRC and associated with carcinogenicity [14, 15, 20, 56]. Therefore, according to these findings, we speculate that the viral genetic integrations in the microbes possibly result in an increasing bacterial diversity and high abundance of some potential pathogens in favor of a dysbiosis, which potentially could be an important factor for carcinogenic process.

**Fig. 5.**
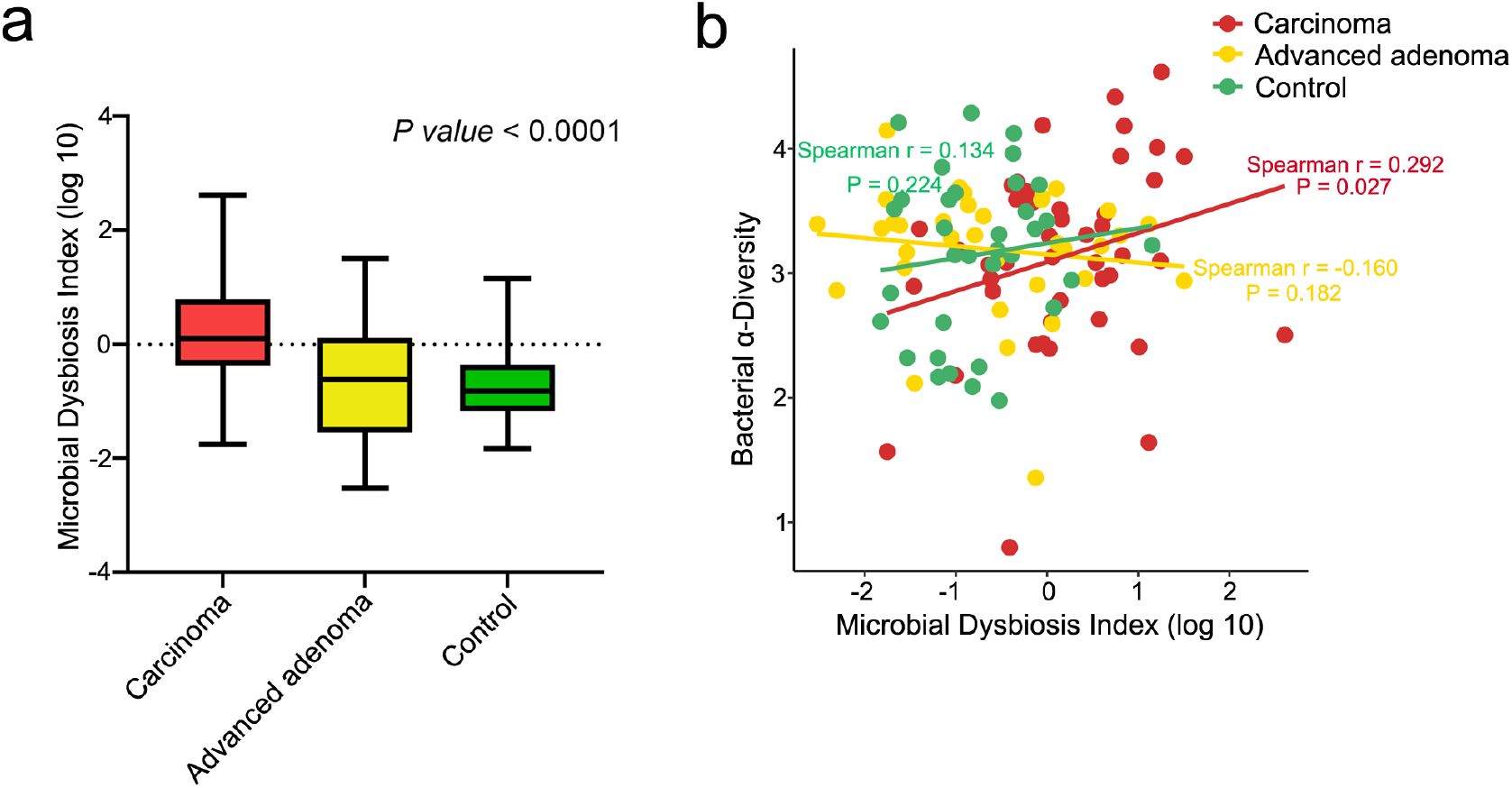
Microbial dysbiosis associated with colorecctal carcinoma. (a). Box plot shows the MDI of the carcinoma, advanced adenoma patients and healthy controls. Significance difference was obtained by one-way analysis of variance (ANOVA) corrected with Kruskal-Wallis test for multiple comparisons. (b). Spearman correlation between MDI and bacterial Shannon index diversity in the different groups. The MDI was positively correlated with its bacterial diversity in carcinoma, while not in advanced adenoma patients and healthy controls. Each dot represents one sample. Healthy controls n = 34, advanced adenoma patients n=34 and carcinoma patients n=44.

### The ubiquitous genetic integration from phage to bacteria in the human gut

Understanding the intricate genetic integration of phages and their bacterial hosts will hopefully help us to further our theories on the microbial interactions. In this part, our purpose is to facilitate researches to select bacteriophages that can interact with a bacteria specifically. Therefore, to ensure the stability of the interaction, the genetic integration of any phages in the bacteria presented in at least twice in different samples were illustrated. As shown, we summarized a distribution of 11 genera of prokaryotic hosts including a range of commensal and potential pathogenic bacteria (Fig. 6). Moreover, we found that almost all of the phages had their hosts at only one genus levels. That is to say, the phages usually have a narrow range of hosts. This result is consistent with the findings of the previous study [57], and further confirms the reliability of data and strict analysis pipeline. Consequently, the genetic integration network will plays a key role in gut microbiome interactions, for instance, the development of phage therapy as an alternative to antibiotics.

**Fig. 6.**
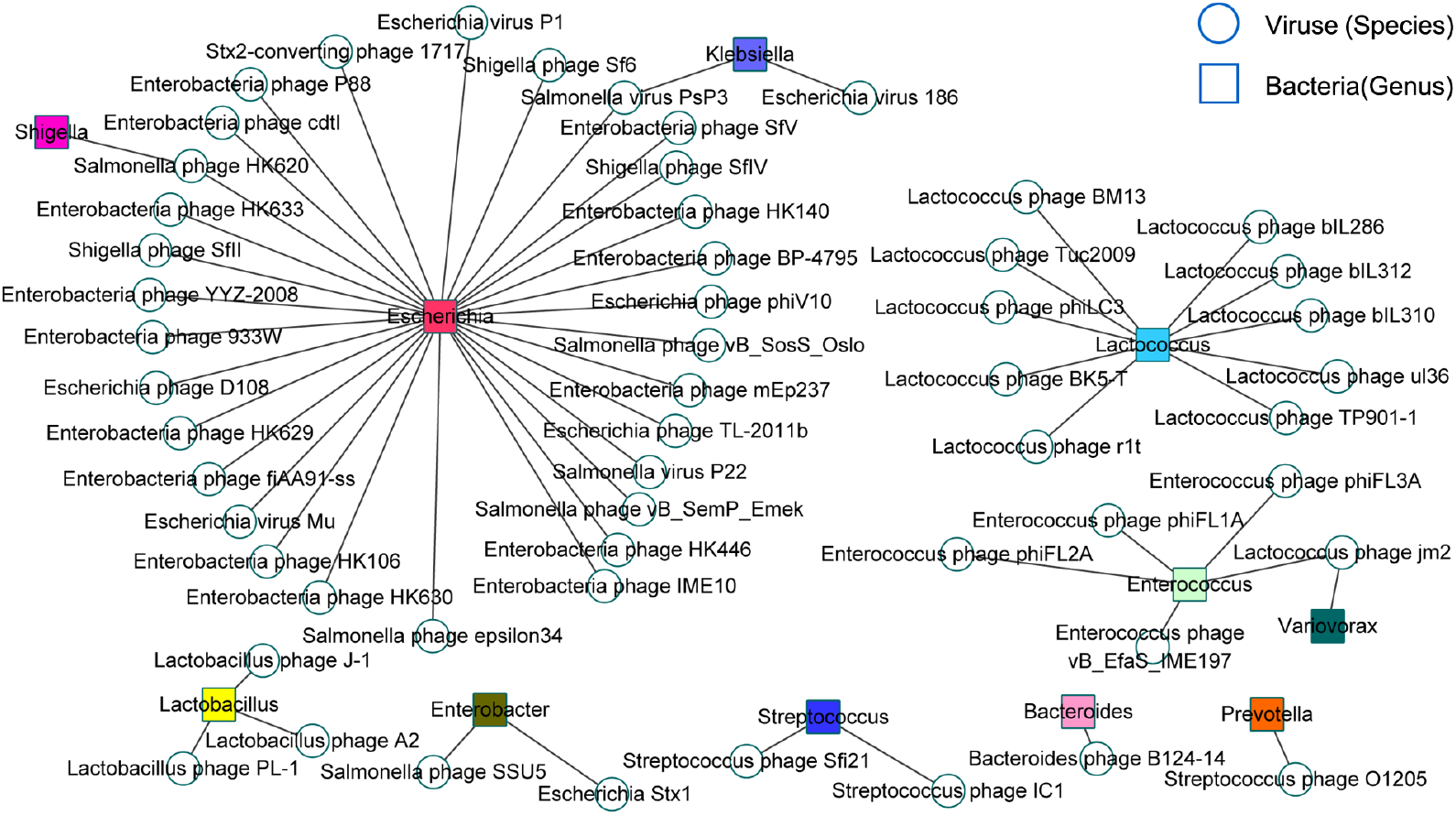
The common phage-bacteria genetic integration network in all of the three cohorts. Within the network, circles represent for phages species and squares for bacteria genera. The darkness of the color nodes indicate their frequency of interactions. The genetic integration network occurred at least twice in different samples was shown.

## Discussion

Analysis of a multi-kingdom gut microbiome interaction contributes to the new research opportunities for understanding the pathogenesis of colorectal carcinoma. In this paper, to our knowledge, we demonstrate for the first time that disease-specific viral genetic integrations in the microbes increase the successful subpopulation of some commensal and potential pathogenic bacteria along with a microbial dysbiosis that potentially could be associated with the occurrence of tumour. We found that the genetic integrations from phages to bacterium mainly appeared in colorectal carcinoma patients and were positively associated with the abundance of the phage communities. Besides, the integration of host-beneficial genes into microbial genomes increased the abundance of commensal and potential pathogenic bacteria. Importantly, the genetic integrations from phages to microbes potentially acted as a factor for the microbial dysbiosis and high bacterial diversity especially in carcinoma patients, which could be an important reason for carcinogenesis of colorectum. In short, we suggest that the viral genetic integrations in microbes contribute to the expansion of some potential pathogens along with the microbial dysbiosis during the human colorectal tumorigenesis.

Undoubtedly, bacteriophages are account for a main portion of virome in the human gut. Previous studies have demonstrated that the abundance of intestinal phages varied with different disease status (i.e., health or carcinoma) [28, 58]. However, the consequences of change on phages in the gut remain poorly known. Our finding identified that the bacteriophages with high abundance had more chances to invade microbes and were able to transfer their own DNA to bacterial genomes. As a result, a large and diverse pool of phages with bacterial-beneficial genes could be inserted into the genomes of bacterial hosts, especially in carcinoma patients, which potentially has a direct role in improving metabolism or cell activity, increasing the subpopulation or pathogenicity of bacteria. Meanwhile, recent studies have shown not only phage from lytic cycle can enter the lysogenic cycle in same case [59], but also the temperate phages can directly insert into the chromosome of the bacteria (called prophages), which could protect their bacterial hosts against secondary infections through superinfection exclusion [30] or CRISPR/Cas immune defense [60]. Based on that, it looks like maybe these are the reasons for the positive correlation between the integration of phages and the abundance of their bacterial hosts in carcinoma patients. Therefore, the specific viral genetic integrations could alter the gut environment in terms of microbial composition, structure, and functionality via selection on characteristics that affect the ability of fighting against competitors and pathogenicity of bacteria.

Now it is generally believed that colorectal carcinoma is closely related to the disorder of intestinal microbial population. As previous studies have demonstrated, the dysbiosis of the gut microbial community as well as gut virome were associated with CRC [24, 58, 61]. And significant difference had been uncovered in bacterial communities between healthy individuals and patients suffered from colorectal cancer by 16S or WGS sequencing [52, 62]. According to our findings, we suggest that the horizontal gene transfer (HGT) from phage to bacteria is likely to play an important role in microbial diversity and multifunction in the human gut of colorectal carcinoma patients. That is to say, the differences of bacteriophage genetic integrations in the microbes between health controls and CRC patients might represent distinct functional roles that are important for microbial functionality and subpopulation, shaping immune system of human. As a result, from point of view of carcinogenic mechanism, the integrated bacteria with altered functionalities (e.g., the new ability to produce or metabolise potential substrates) provides an opportunity for establishment of dysbiosis ecological networks within the gut microbial population, which is possibly associated with the development of tumor microbiome, modulate the immune system and, ultimately, affect tumor growth.

Viruses and bacteria are key components of the human microbiome and previous works on human microbiome have mainly focused on bacterial and viral independently [27]. Thus these strategies fail to capture the complex dynamics of interaction between bacteria and phage. Studying such phage-bacteria integrations contribute us to discovery the complex interaction in the human gut and provides the details networks of phages and bacterium infection in different disease status. The identification that who infects whom is essential to enhance our theories on the intricacies of these microbial interactions and to understand the functional influence of infection in complex ecosystem such as changing metabolic products and determining the fate of hosts. Moreover, with increasing severity of the drug-resistant superbacteria, the phage-bacteria networks offer the theoretical foundation to use phages in treatment of pathogenic bacteria. Therefore, this work will plays a pivotal role in the development of phage therapy as an alternative to antibiotics.

In summary, our metagenomic association survey on the interaction of virus-bacteria was performed in order to identify the genetic integrations from viruses to microbial communities and their association with the colorectal tumorigenesis. The integration of phage-carried positive effective genes were able to cause the increasing abundance of some bacterium including potential pathogens along with microbial dysbiosis in carcinoma patients. These results imply that phages potentially play an important role in morphogenesis of gut microbial community and consequently affect the healthy status. Besides, the genetic integration networks could be of a great use in the treatment of drug-resistant bacteria by phage therapy. Therefore, we suggest a potential role that utilization of phage on therapy of colorectal diseases. However, additional work will be imperative to further validate the conclusion in more large cohorts or by experimental study.

## Acknowledgments

We appreciate the helpful discussion with Dr. Kunming Li. This work was supported by grants from the National Key R&D Program of China [2018YFC0910402 to J.C.; 2018YFC1003102 to C.Z. and 2017YFC0908402 to C.Z.]; the National Natural Science Foundation of China [31571307 to C.J.]; the Open Project of Key Laboratory of Genomic and Precision Medicine, Chinese Academy of Sciences.

## Authors’ contributions

BC and XW performed all the analyses. JC, BC and CZ designed the study. JC, BC and WX wrote the manuscript. All authors read and approved the final manuscript.

## Ethics approval and consent to participate

Not applicable.

## Supplemental figure and table legend

**Fig. S1.**
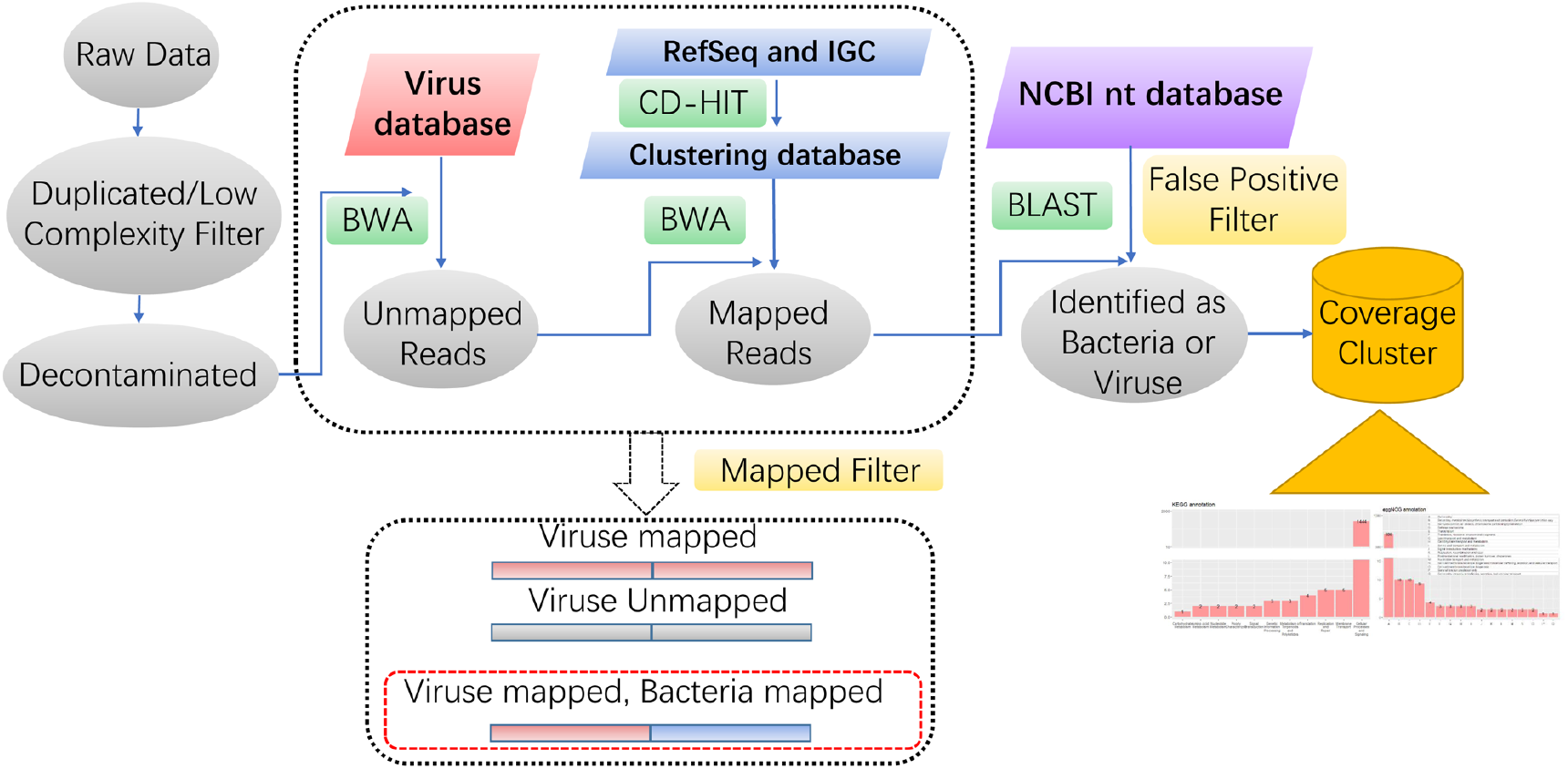
Detailed schematic of computational analysis workflow to identify putative genetic integration events.

**Fig. S2.**
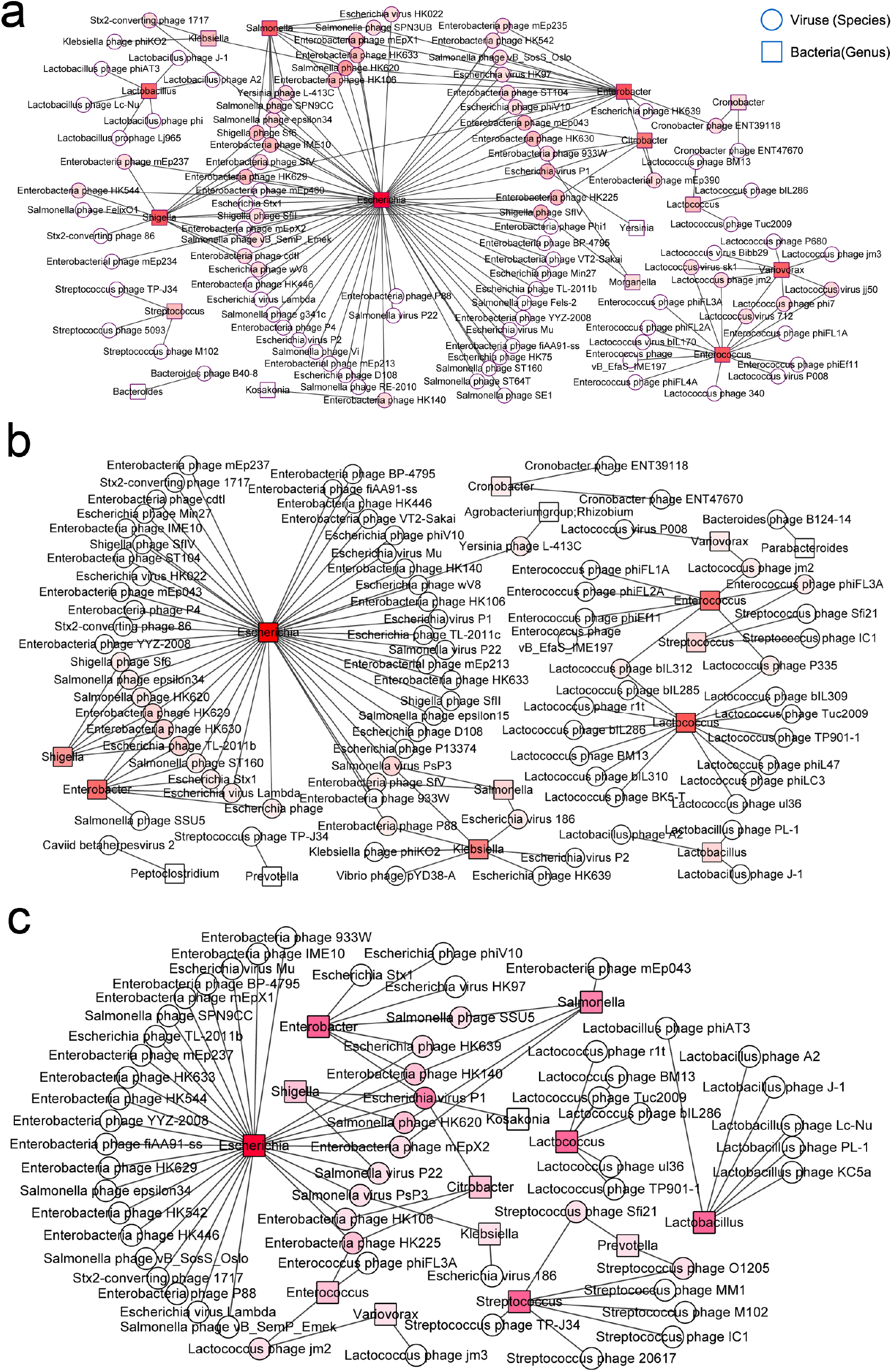
The phage-bacteria genetic integration network in the human gut of three groups. Within the network, phages species are represented as circle nodes, while the bacterial genera displayed as square nodes. The higher frequency are shown as darker colour nodes. The viral genetic integrations in microbes were more prevalent in carcinoma than in healthy controls, follow by advanced adenoma patients.

**Fig. S3.**
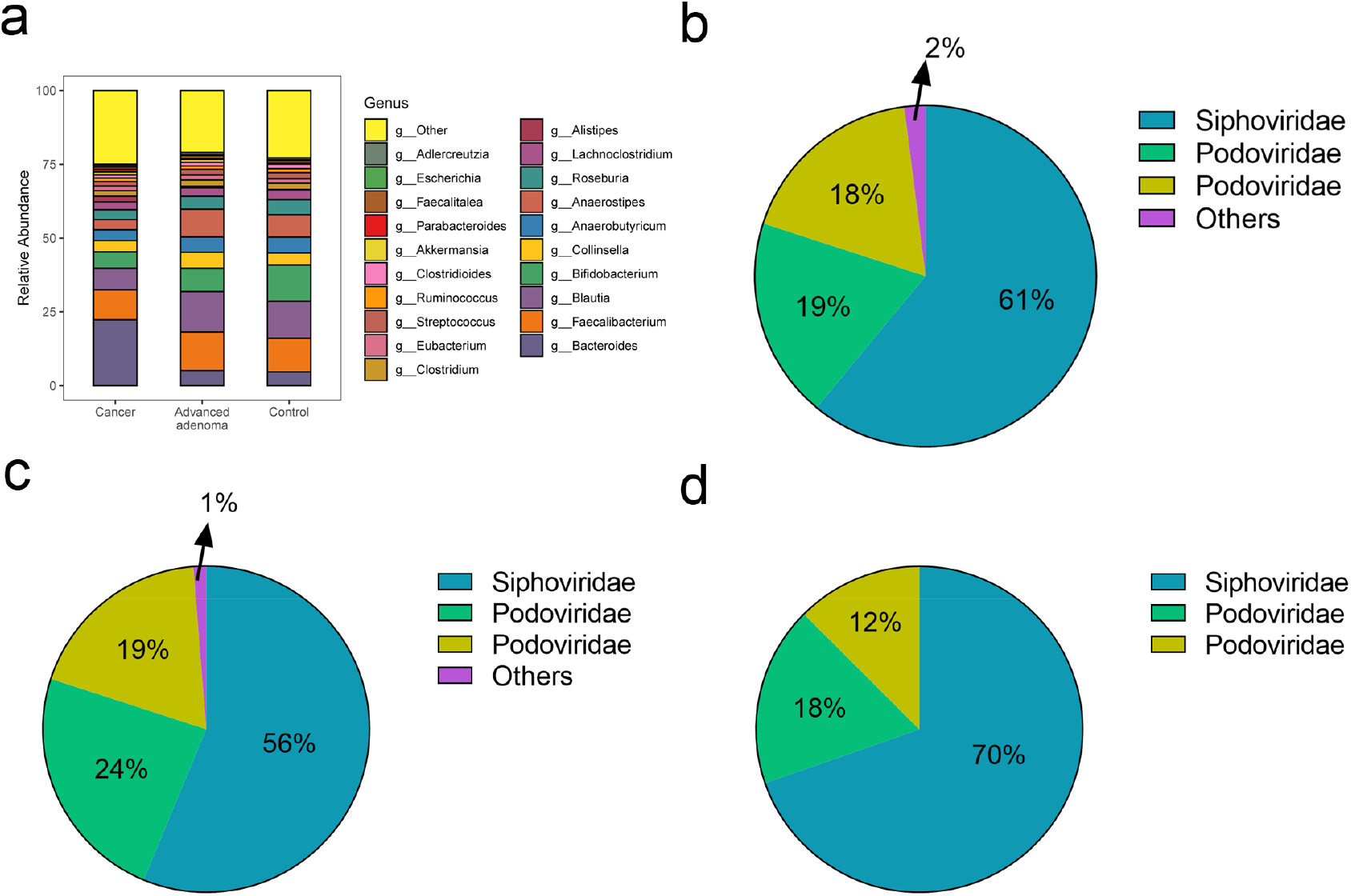
The relative abundance of bacterial genera in the three groups. The Difference of gut microbial communities were shown in carcinoma, advanced adenoma patients and healthy control groups (a). The families of viral genetic integrations into their hosts were shown in carcinoma (b), advanced adenoma patients (c) and healthy control groups (d).

**Fig. S4.**
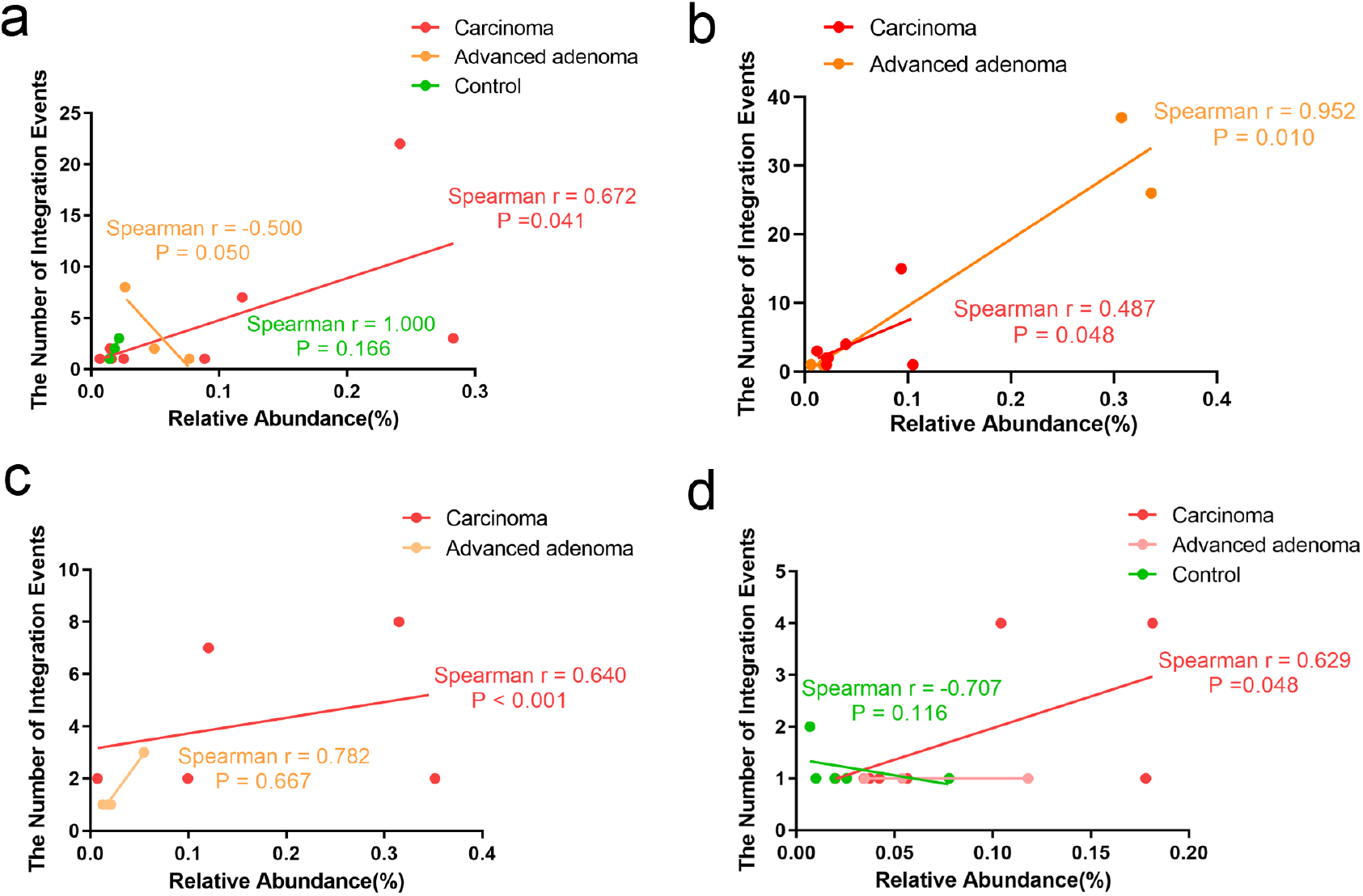
The relationship between the integrated frequency of *Enterobacter* (a), *Enterococcus* (b), *Shigella* (c) and *Bacteroides* (d) and their relative abundance, respectively. The genetic integration of the bacterial genera were positively correlated with their abundance in carcinoma patients, while not in healthy control and advanced adenoma patients. Each dot represents one sample and the Spearman correlation coefficients are indicated.

**Fig. S5.**
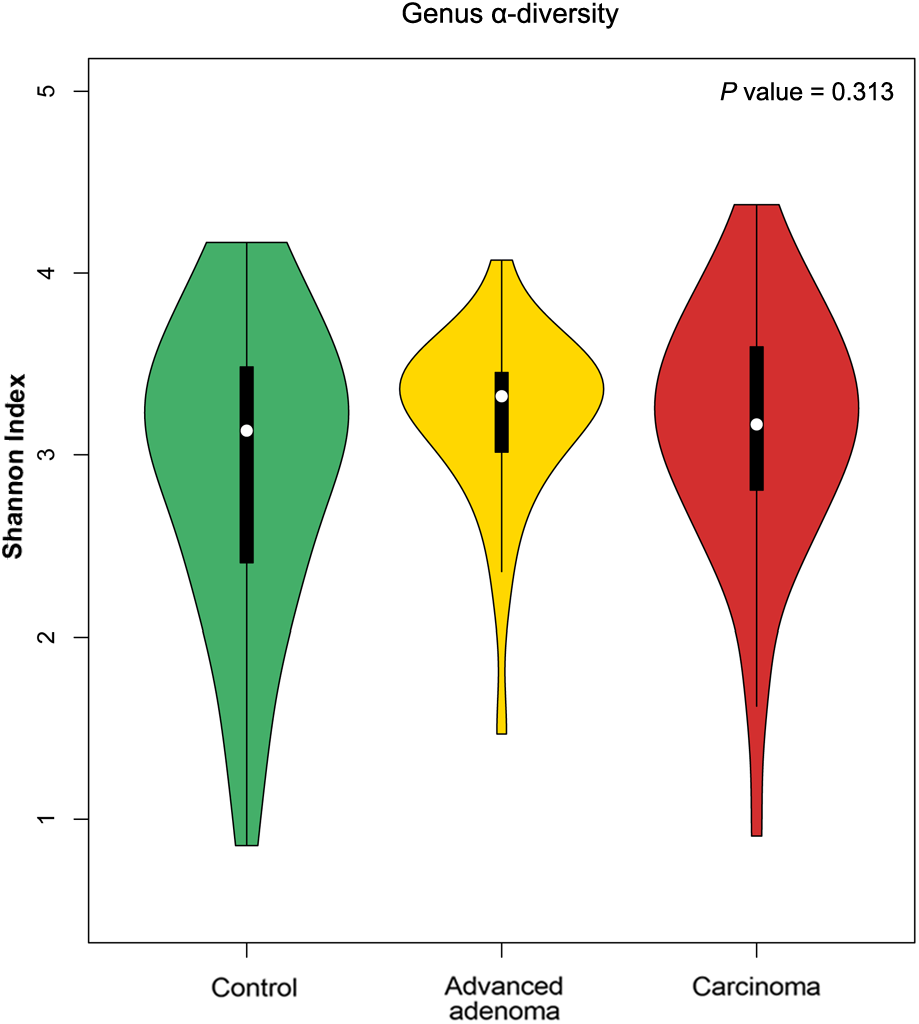
The bacterial a-diversity (Shannon index) of the three groups at the genus level. *P value* from Kruskal-Wallis test is shown.

**Table S1.** The information of samples used in this study.

**Table S2**. The phage taxonomy of the genetic integration network.

## References

1. Colorectal cancer: new evidence for the adenoma/carcinoma sequence. Lancet. 1992;340(8813):210–1. Epub 1992/07/25. PubMed PMID: 1353140.

2. Leslie A, Carey FA, Pratt NR, Steele RJ. The colorectal adenoma-carcinoma sequence. Br J Surg. 2002;89(7):845–60. Epub 2002/06/26. doi: 10.1046/j.1365-2168.2002.02120.x. PubMed PMID: 12081733.

3. Maesawa C, Tamura G, Suzuki Y, Ogasawara S, Sakata K, Kashiwaba M, et al. The sequential accumulation of genetic alterations characteristic of the colorectal adenoma-carcinoma sequence does not occur between gastric adenoma and adenocarcinoma. J Pathol. 1995;176(3):249–58. Epub 1995/07/01. doi: 10.1002/path.1711760307. PubMed PMID: 7674088.

4. Feng Q, Liang S, Jia H, Stadlmayr A, Tang L, Lan Z, et al. Gut microbiome development along the colorectal adenoma-carcinoma sequence. Nat Commun. 2015;6:6528. Epub 2015/03/12. doi: 10.1038/ncomms7528. PubMed PMID: 25758642.

5. Ferreira RM, Pereira-Marques J, Pinto-Ribeiro I, Costa JL, Carneiro F, Machado JC, et al. Gastric microbial community profiling reveals a dysbiotic cancer-associated microbiota. Gut. 2018;67(2):226–36. Epub 2017/11/06. doi: 10.1136/gutjnl-2017-314205. PubMed PMID: 29102920; PubMed Central PMCID: PMCPMC5868293.

6. Thomas AM, Manghi P, Asnicar F, Pasolli E, Armanini F, Zolfo M, et al. Metagenomic analysis of colorectal cancer datasets identifies cross-cohort microbial diagnostic signatures and a link with choline degradation. Nat Med. 2019;25(4):667–78. Epub 2019/04/03. doi: 10.1038/s41591-019-0405-7. PubMed PMID: 30936548.

7. Sender R, Fuchs S, Milo R. Revised Estimates for the Number of Human and Bacteria Cells in the Body. PLoS Biol. 2016;14(8):e1002533. Epub 2016/08/20. doi: 10.1371/journal.pbio.1002533. PubMed PMID: 27541692; PubMed Central PMCID: PMCPMC4991899.

8. Breitbart M, Hewson I, Felts B, Mahaffy JM, Nulton J, Salamon P, et al. Metagenomic analyses of an uncultured viral community from human feces. J Bacteriol. 2003;185(20):6220–3. Epub 2003/10/04. doi: 10.1128/jb.185.20.6220-6223.2003. PubMed PMID: 14526037; PubMed Central PMCID: PMCPMC225035.

9. Minot S, Sinha R, Chen J, Li H, Keilbaugh SA, Wu GD, et al. The human gut virome: interindividual variation and dynamic response to diet. Genome Res. 2011;21(10):1616–25. Epub 2011/09/02. doi: 10.1101/gr.122705.111. PubMed PMID: 21880779; PubMed Central PMCID: PMCPMC3202279.

10. Reyes A, Haynes M, Hanson N, Angly FE, Heath AC, Rohwer F, et al. Viruses in the faecal microbiota of monozygotic twins and their mothers. Nature. 2010;466(7304):334–8. Epub 2010/07/16. doi: 10.1038/nature09199. PubMed PMID: 20631792; PubMed Central PMCID: PMCPMC2919852.

11. Zhu Q, Gao R, Wu W, Qin H. The role of gut microbiota in the pathogenesis of colorectal cancer. Tumour Biol. 2013;34(3):1285–300. Epub 2013/02/12. doi: 10.1007/s13277-013-0684-4. PubMed PMID: 23397545.

12. Nakatsu G, Li X, Zhou H, Sheng J, Wong SH, Wu WK, et al. Gut mucosal microbiome across stages of colorectal carcinogenesis. Nat Commun. 2015;6:8727. Epub 2015/10/31. doi: 10.1038/ncomms9727. PubMed PMID: 26515465; PubMed Central PMCID: PMCPMC4640069.

13. Mima K, Ogino S, Nakagawa S, Sawayama H, Kinoshita K, Krashima R, et al. The role of intestinal bacteria in the development and progression of gastrointestinal tract neoplasms. Surg Oncol. 2017;26(4):368–76. Epub 2017/11/09. doi: 10.1016/j.suronc.2017.07.011. PubMed PMID: 29113654; PubMed Central PMCID: PMCPMC5726560.

14. Castellarin M, Warren RL, Freeman JD, Dreolini L, Krzywinski M, Strauss J, et al. Fusobacterium nucleatum infection is prevalent in human colorectal carcinoma. Genome Res. 2012;22(2):299–306. Epub 2011/10/20. doi: 10.1101/gr.126516.111. PubMed PMID: 22009989; PubMed Central PMCID: PMCPMC3266037.

15. Gagniere J, Raisch J, Veziant J, Barnich N, Bonnet R, Buc E, et al. Gut microbiota imbalance and colorectal cancer. World J Gastroenterol. 2016;22(2):501–18. Epub 2016/01/27. doi: 10.3748/wjg.v22.i2.501. PubMed PMID: 26811603; PubMed Central PMCID: PMCPMC4716055.

16. Louis P, Hold GL, Flint HJ. The gut microbiota, bacterial metabolites and colorectal cancer. Nat Rev Microbiol. 2014;12(10):661–72. Epub 2014/09/10. doi: 10.1038/nrmicro3344. PubMed PMID: 25198138.

17. Coleman OI, Haller D. Bacterial Signaling at the intestinal epithelial interface in inflammation and Cancer. Frontiers in Immunology. 2018;8. PubMed PMID: WOS:000419376000001.

18. Cremonesi E, Governa V, Garzon JFG, Mele V, Amicarella F, Muraro MG, et al. Gut microbiota modulate T cell trafficking into human colorectal cancer. Gut. 2018;67(11):1984–94. PubMed PMID: WOS:000448519800012.

19. Bashir A, Miskeen AY, Hazari YM, Asrafuzzaman S, Fazili KM. Fusobacterium nucleatum, inflammation, and immunity: the fire within human gut. Tumor Biol. 2016;37(3):2805–10. PubMed PMID: WOS:000374903500003.

20. Zou SM, Fang LK, Lee MH. Dysbiosis of gut microbiota in promoting the development of colorectal cancer. Gastroenterol Rep. 2018;6(1):1–12. PubMed PMID: WOS:000429991500001.

21. Richard ML, Liguori G, Lamas B, Brandi G, da Costa G, Hoffmann TW, et al. Mucosa-associated microbiota dysbiosis in colitis associated cancer. Gut Microbes. 2018;9(2):131–42. PubMed PMID: WOS:000435713000005.

22. Gao ZG, Guo BM, Gao RY, Zhu QC, Qin HL. Microbiota disbiosis is associated with colorectal cancer. Front Microbiol. 2015;6. PubMed PMID: WOS:000349999700001.

23. Gao Z, Guo B, Gao R, Zhu Q, Qin H. Microbiota disbiosis is associated with colorectal cancer. Front Microbiol. 2015;6:20. Epub 2015/02/24. doi: 10.3389/fmicb.2015.00020. PubMed PMID: 25699023; PubMed Central PMCID: PMCPMC4313696.

24. Wu N, Yang X, Zhang RF, Li J, Xiao X, Hu YF, et al. Dysbiosis Signature of Fecal Microbiota in Colorectal Cancer Patients. Microbial Ecology. 2013;66(2):462–70. PubMed PMID: WOS:000321668900020.

25. Akin H, Tozun N. Diet, Microbiota, and Colorectal Cancer. Journal of clinical gastroenterology. 2014;48:S67–S9. PubMed PMID: WOS:000347246300017.

26. Goodrich JK, Waters JL, Poole AC, Sutter JL, Koren O, Blekhman R, et al. Human genetics shape the gut microbiome. Cell. 2014;159(4):789–99. Epub 2014/11/25. doi: 10.1016/j.cell.2014.09.053. PubMed PMID: 25417156; PubMed Central PMCID: PMCPMC4255478.

27. Hannigan GD, Duhaime MB, Koutra D, Schloss PD. Biogeography and environmental conditions shape bacteriophage-bacteria networks across the human microbiome. PLoS Comput Biol. 2018;14(4):e1006099. Epub 2018/04/19. doi: 10.1371/journal.pcbi.1006099. PubMed PMID: 29668682; PubMed Central PMCID: PMCPMC5927471.

28. Nakatsu G, Zhou H, Wu WKK, Wong SH, Coker OO, Dai Z, et al. Alterations in Enteric Virome Are Associated With Colorectal Cancer and Survival Outcomes. Gastroenterology. 2018;155(2):529–41 e5. Epub 2018/04/25. doi: 10.1053/j.gastro.2018.04.018. PubMed PMID: 29689266.

29. Brown SP, Le Chat L, De Paepe M, Taddei F. Ecology of microbial invasions: amplification allows virus carriers to invade more rapidly when rare. Curr Biol. 2006;16(20):2048–52. Epub 2006/10/24. doi: 10.1016/j.cub.2006.08.089. PubMed PMID: 17055985.

30. Obeng N, Pratama AA, Elsas JDV. The Significance of Mutualistic Phages for Bacterial Ecology and Evolution. Trends Microbiol. 2016;24(6):440–9. Epub 2016/02/02. doi: 10.1016/j.tim.2015.12.009. PubMed PMID: 26826796.

31. Hargreaves KR, Kropinski AM, Clokie MR. Bacteriophage behavioral ecology: How phages alter their bacterial host’s habits. Bacteriophage. 2014;4:e29866. Epub 2014/08/12. doi: 10.4161/bact.29866. PubMed PMID: 25105060; PubMed Central PMCID: PMCPMC4124054.

32. Rodriguez-Valera F, Martin-Cuadrado AB, Rodriguez-Brito B, Pasic L, Thingstad TF, Rohwer F, et al. Explaining microbial population genomics through phage predation. Nat Rev Microbiol. 2009;7(11):828–36. Epub 2009/10/17. doi: 10.1038/nrmicro2235. PubMed PMID: 19834481.

33. Miao EA, Miller SI. Bacteriophages in the evolution of pathogen-host interactions. Proc Natl Acad Sci U S A. 1999;96(17):9452–4. Epub 1999/08/18. doi: 10.1073/pnas.96.17.9452. PubMed PMID: 10449711; PubMed Central PMCID: PMCPMC33707.

34. Bolger AM, Lohse M, Usadel B. Trimmomatic: a flexible trimmer for Illumina sequence data. Bioinformatics. 2014;30(15):2114–20. Epub 2014/04/04. doi: 10.1093/bioinformatics/btu170. PubMed PMID: 24695404; PubMed Central PMCID: PMCPMC4103590.

35. Li H, Handsaker B, Wysoker A, Fennell T, Ruan J, Homer N, et al. The Sequence Alignment/Map format and SAMtools. Bioinformatics. 2009;25(16):2078–9. Epub 2009/06/10. doi: 10.1093/bioinformatics/btp352. PubMed PMID: 19505943; PubMed Central PMCID: PMCPMC2723002.

36. Li J, Jia H, Cai X, Zhong H, Feng Q, Sunagawa S, et al. An integrated catalog of reference genes in the human gut microbiome. Nat Biotechnol. 2014;32(8):834–41. Epub 2014/07/07. doi: 10.1038/nbt.2942. PubMed PMID: 24997786.

37. Camacho C, Coulouris G, Avagyan V, Ma N, Papadopoulos J, Bealer K, et al. BLAST+: architecture and applications. BMC Bioinformatics. 2009;10:421. doi: 10.1186/1471-2105-10-421. PubMed PMID: 20003500; PubMed Central PMCID: PMC2803857.

38. Riley DR, Sieber KB, Robinson KM, White JR, Ganesan A, Nourbakhsh S, et al. Bacteria-human somatic cell lateral gene transfer is enriched in cancer samples. PLoS Comput Biol. 2013;9(6):e1003107. Epub 2013/07/11. doi: 10.1371/journal.pcbi.1003107. PubMed PMID: 23840181; PubMed Central PMCID: PMCPMC3688693.

39. Wood DE, Salzberg SL. Kraken: ultrafast metagenomic sequence classification using exact alignments. Genome Biol. 2014;15(3):R46. Epub 2014/03/04. doi: 10.1186/gb-2014-15-3-r46. PubMed PMID: 24580807; PubMed Central PMCID: PMCPMC4053813.

40. Dixon P. VEGAN, a package of R functions for community ecology. J Veg Sci. 2003;14(6):927–30. doi: Doi 10.1658/1100-9233(2003)014[0927:Vaporf]2.0.Co;2. PubMed PMID: WOS:000189220100019.

41. Huerta-Cepas J, Szklarczyk D, Forslund K, Cook H, Heller D, Walter MC, et al. eggNOG 4.5: a hierarchical orthology framework with improved functional annotations for eukaryotic, prokaryotic and viral sequences. Nucleic Acids Res. 2016;44(D1):D286–93. Epub 2015/11/20. doi: 10.1093/nar/gkv1248. PubMed PMID: 26582926; PubMed Central PMCID: PMCPMC4702882.

42. Otasek D, Morris JH, Boucas J, Pico AR, Demchak B. Cytoscape Automation: empowering workflow-based network analysis. Genome Biol. 2019;20(1):185. Epub 2019/09/04. doi: 10.1186/s13059-019-1758-4. PubMed PMID: 31477170; PubMed Central PMCID: PMCPMC6717989.

43. Vogelmann R, Amieva MR. The role of bacterial pathogens in cancer. Curr Opin Microbiol. 2007;10(1):76–81. Epub 2007/01/09. doi: 10.1016/j.mib.2006.12.004. PubMed PMID: 17208515.

44. Leung A, Tsoi H, Yu J. Fusobacterium and Escherichia: models of colorectal cancer driven by microbiota and the utility of microbiota in colorectal cancer screening. Expert Rev Gastroenterol Hepatol. 2015;9(5):651–7. Epub 2015/01/15. doi: 10.1586/17474124.2015.1001745. PubMed PMID: 25582922.

45. Zhang N, He QS. Commensal Microbiome Promotes Resistance to Local and Systemic Infections. Chin Med J (Engl). 2015;128(16):2250–5. Epub 2015/08/13. doi: 10.4103/0366-6999.162502. PubMed PMID: 26265621; PubMed Central PMCID: PMCPMC4717980.

46. Arleevskaya MI, Aminov R, Brooks WH, Manukyan G, Renaudineau Y. Editorial: Shaping of Human Immune System and Metabolic Processes by Viruses and Microorganisms. Front Microbiol. 2019;10:816. Epub 2019/05/07. doi: 10.3389/fmicb.2019.00816. PubMed PMID: 31057521; PubMed Central PMCID: PMCPMC6481243.

47. Moreno-Gallego JL, Chou SP, Di Rienzi SC, Goodrich JK, Spector TD, Bell JT, et al. Virome Diversity Correlates with Intestinal Microbiome Diversity in Adult Monozygotic Twins. Cell Host Microbe. 2019;25(2):261–72 e5. Epub 2019/02/15. doi: 10.1016/j.chom.2019.01.019. PubMed PMID: 30763537; PubMed Central PMCID: PMCPMC6411085.

48. Huerta-Cepas J, Forslund K, Coelho LP, Szklarczyk D, Jensen LJ, von Mering C, et al. Fast Genome-Wide Functional Annotation through Orthology Assignment by eggNOG-Mapper. Mol Biol Evol. 2017;34(8):2115–22. Epub 2017/05/02. doi: 10.1093/molbev/msx148. PubMed PMID: 28460117; PubMed Central PMCID: PMCPMC5850834.

49. Gevers D, Kugathasan S, Denson LA, Vazquez-Baeza Y, Van Treuren W, Ren B, et al. The treatment-naive microbiome in new-onset Crohn’s disease. Cell Host Microbe. 2014;15(3):382–92. Epub 2014/03/19. doi: 10.1016/j.chom.2014.02.005. PubMed PMID: 24629344; PubMed Central PMCID: PMCPMC4059512.

50. Zhu QC, Gao RY, Wu W, Qin HL. The role of gut microbiota in the pathogenesis of colorectal cancer. Tumor Biol. 2013;34(3):1285–300. doi: 10.1007/s13277-013-0684-4. PubMed PMID: WOS:000319356900002.

51. Dai Z, Coker OO, Nakatsu G, Wu WKK, Zhao L, Chen Z, et al. Multi-cohort analysis of colorectal cancer metagenome identified altered bacteria across populations and universal bacterial markers. Microbiome. 2018;6(1):70. Epub 2018/04/13. doi: 10.1186/s40168-018-0451-2. PubMed PMID: 29642940; PubMed Central PMCID: PMCPMC5896039.

52. Yu J, Feng Q, Wong SH, Zhang D, Liang QY, Qin Y, et al. Metagenomic analysis of faecal microbiome as a tool towards targeted non-invasive biomarkers for colorectal cancer. Gut. 2017;66(1):70–8. Epub 2015/09/27. doi: 10.1136/gutjnl-2015-309800. PubMed PMID: 26408641.

53. Wirbel J, Pyl PT, Kartal E, Zych K, Kashani A, Milanese A, et al. Meta-analysis of fecal metagenomes reveals global microbial signatures that are specific for colorectal cancer. Nat Med. 2019;25(4):679–89. Epub 2019/04/03. doi: 10.1038/s41591-019-0406-6. PubMed PMID: 30936547.

54. Gao R, Gao Z, Huang L, Qin H. Gut microbiota and colorectal cancer. Eur J Clin Microbiol Infect Dis. 2017;36(5):757–69. Epub 2017/01/08. doi: 10.1007/s10096-016-2881-8. PubMed PMID: 28063002; PubMed Central PMCID: PMCPMC5395603.

55. Wang T, Cai G, Qiu Y, Fei N, Zhang M, Pang X, et al. Structural segregation of gut microbiota between colorectal cancer patients and healthy volunteers. ISME J. 2012;6(2):320–9. Epub 2011/08/19. doi: 10.1038/ismej.2011.109. PubMed PMID: 21850056; PubMed Central PMCID: PMCPMC3260502.

56. Kostic AD, Chun EY, Robertson L, Glickman JN, Gallini CA, Michaud M, et al. Fusobacterium nucleatum Potentiates Intestinal Tumorigenesis and Modulates the Tumor-Immune Microenvironment. Cell Host & Microbe. 2013;14(2):207–15. PubMed PMID: WOS:000330851600011.

57. Wang J, Gao Y, Zhao F. Phage-bacteria interaction network in human oral microbiome. Environ Microbiol. 2016;18(7):2143–58. Epub 2015/06/04. doi: 10.1111/1462-2920.12923. PubMed PMID: 26036920.

58. Hannigan GD, Duhaime MB, Ruffin MTt, Koumpouras CC, Schloss PD. Diagnostic Potential and Interactive Dynamics of the Colorectal Cancer Virome. MBio. 2018;9(6). Epub 2018/11/22. doi: 10.1128/mBio.02248-18. PubMed PMID: 30459201; PubMed Central PMCID: PMCPMC6247079.

59. Maurice CF, Bouvier C, de Wit R, Bouvier T. Linking the lytic and lysogenic bacteriophage cycles to environmental conditions, host physiology and their variability in coastal lagoons. Environ Microbiol. 2013;15(9):2463–75. Epub 2013/04/16. doi: 10.1111/1462-2920.12120. PubMed PMID: 23581698.

60. Deveau H, Garneau JE, Moineau S. CRISPR/Cas system and its role in phage-bacteria interactions. Annu Rev Microbiol. 2010;64:475–93. Epub 2010/06/10. doi: 10.1146/annurev.micro.112408.134123. PubMed PMID: 20528693.

61. Zou S, Fang L, Lee MH. Dysbiosis of gut microbiota in promoting the development of colorectal cancer. Gastroenterol Rep (Oxf). 2018;6(1):1–12. Epub 2018/02/27. doi: 10.1093/gastro/gox031. PubMed PMID: 29479437; PubMed Central PMCID: PMCPMC5806407.

62. Osman MA, Neoh HM, Ab Mutalib NS, Chin SF, Jamal R. 16S rRNA Gene Sequencing for Deciphering the Colorectal Cancer Gut Microbiome: Current Protocols and Workflows. Front Microbiol. 2018;9. PubMed PMID: WOS:000431017300002.

