## Supplementary material for "The virus integrations in gut microbes associated with dysbiosis of microbial community in tumorigenesis of colorectal carcinoma": Table S1

**Table S1. The information of samples used in this study.**

| Sample ID | State | Age (yrs) | Gender (1:male, 2:female) | Run_accession |
|---|---|---|---|---|
| 31071 | controls | 68 | 1 | ERR688509 |
| 31112 | controls | 66 | 1 | ERR688510 |
| 31129 | controls | 73 | 1 | ERR688511 |
| 31137 | advanced adenoma | 67 | 1 | ERR688512 |
| 31159 | carcinoma | 73 | 1 | ERR688513 |
| 31160 | controls | 67 | 2 | ERR688514 |
| 31188 | carcinoma | 65 | 1 | ERR688515 |
| 31219 | controls | 74 | 2 | ERR688516 |
| 31223 | carcinoma | 65 | 2 | ERR688517 |
| 31232 | controls | 70 | 1 | ERR688518 |
| 31233 | advanced adenoma | 62 | 1 | ERR688519 |
| 31236 | controls | 65 | 1 | ERR688520 |
| 31237 | carcinoma | 67 | 1 | ERR688521 |
| 31256 | advanced adenoma | 69 | 2 | ERR688522 |
| 31267 | controls | 67 | 1 | ERR688523 |
| 31275 | advanced adenoma | 62 | 1 | ERR688524 |
| 31276 | carcinoma | 82 | 1 | ERR688525 |
| 31282 | advanced adenoma | 63 | 1 | ERR688526 |
| 31285 | carcinoma | 84 | 1 | ERR688527 |
| 31300 | controls | 68 | 2 | ERR688528 |
| 31328 | controls | 63 | 1 | ERR688529 |
| 31333 | controls | 68 | 1 | ERR688530 |
| 31337 | advanced adenoma | 62 | 1 | ERR688531 |
| 31343 | controls | 72 | 1 | ERR688532 |
| 31360 | controls | 71 | 1 | ERR688533 |
| 31367 | carcinoma | 86 | 2 | ERR688534 |
| 31379 | controls | 63 | 1 | ERR688535 |
| 31398 | advanced adenoma | 68 | 2 | ERR688536 |
| 31416 | controls | 72 | 1 | ERR688537 |
| 31424 | advanced adenoma | 68 | 2 | ERR688538 |
| 31428 | controls | 69 | 2 | ERR688539 |
| 31431 | advanced adenoma | 71 | 1 | ERR688540 |
| 31446 | carcinoma | 84 | 2 | ERR688541 |
| 31449 | advanced adenoma | 63 | 1 | ERR688542 |
| 31450 | controls | 67 | 1 | ERR688543 |
| 31452 | controls | 66 | 2 | ERR688544 |
| 31455 | advanced adenoma | 72 | 2 | ERR688545 |
| 31477 | advanced adenoma | 78 | 2 | ERR688546 |
| 31489 | carcinoma | 60 | 1 | ERR688547 |
| 31493 | carcinoma | 68 | 1 | ERR688548 |
| 31501 | advanced adenoma | 69 | 2 | ERR688549 |
| 31512 | controls | 64 | 1 | ERR688550 |
| 31519 | controls | 65 | 1 | ERR688551 |
| 31537 | controls | 70 | 2 | ERR688552 |
| 31549 | carcinoma | 84 | 2 | ERR688553 |
| 31557 | controls | 74 | 2 | ERR688554 |
| 31582 | advanced adenoma | 77 | 1 | ERR688555 |
| 31600 | controls | 70 | 2 | ERR688556 |

| | | | | |
|---|---|---|---|---|
| 31637 | controls | 66 | 1 | ERR688557 |
| 31685 | carcinoma | 74 | 1 | ERR688558 |
| 31700 | controls | 72 | 2 | ERR688559 |
| 31705 | advanced adenoma | 84 | 2 | ERR688560 |
| 31711 | controls | 66 | 1 | ERR688561 |
| 31714 | controls | 69 | 2 | ERR688562 |
| 31723 | controls | 69 | 2 | ERR688563 |
| 31749 | controls | 70 | 1 | ERR688564 |
| 31750 | controls | 66 | 1 | ERR688565 |
| 31766 | controls | 73 | 2 | ERR688566 |
| 31865 | carcinoma | 55 | 2 | ERR688567 |
| 31866 | carcinoma | 73 | 2 | ERR688568 |
| 31868 | carcinoma | 64 | 2 | ERR688569 |
| 31870 | carcinoma | 63 | 1 | ERR688570 |
| 31871 | carcinoma | 72 | 1 | ERR688571 |
| 31872 | carcinoma | 46 | 1 | ERR688572 |
| 31873 | carcinoma | 73 | 1 | ERR688573 |
| 31874 | carcinoma | 74 | 2 | ERR688574 |
| 31875 | carcinoma | 58 | 1 | ERR688575 |
| 31876 | carcinoma | 56 | 2 | ERR688576 |
| 31877 | carcinoma | 72 | 2 | ERR688577 |
| 31878 | carcinoma | 52 | 2 | ERR688578 |
| 31879 | carcinoma | 71 | 1 | ERR688579 |
| 31880 | carcinoma | 74 | 1 | ERR688580 |
| 31881 | carcinoma | 60 | 2 | ERR688581 |
| 31883 | carcinoma | 43 | 1 | ERR688582 |
| 31884 | carcinoma | 54 | 1 | ERR688583 |
| 530002 | advanced adenoma | 60 | 2 | ERR688584 |
| 530018 | advanced adenoma | 56 | 1 | ERR688585 |
| 530026 | advanced adenoma | 83 | 2 | ERR688586 |
| 530028 | advanced adenoma | 67 | 1 | ERR688587 |
| 530168 | advanced adenoma | 62 | 2 | ERR688601 |
| 530172 | advanced adenoma | 63 | 1 | ERR688602 |
| 530185 | advanced adenoma | 52 | 2 | ERR688604 |
| 530215 | controls | 65 | 1 | ERR688605 |
| 530251 | controls | 72 | 1 | ERR688607 |
| 530262 | advanced adenoma | 64 | 1 | ERR688609 |
| 530297 | advanced adenoma | 78 | 2 | ERR688611 |
| 530323 | advanced adenoma | 71 | 1 | ERR688613 |
| 530348 | advanced adenoma | 58 | 1 | ERR688615 |
| 530373 | carcinoma | 47 | 2 | ERR688618 |
| 530398 | advanced adenoma | 80 | 1 | ERR688620 |
| 530403 | advanced adenoma | 75 | 2 | ERR688621 |
| 530450 | advanced adenoma | 48 | 1 | ERR688623 |
| 530558 | advanced adenoma | 56 | 2 | ERR688625 |
| 530600 | advanced adenoma | 70 | 2 | ERR688626 |
| 530623 | advanced adenoma | 62 | 1 | ERR688627 |
| 530697 | advanced adenoma | 63 | 2 | ERR688628 |
| 530705 | advanced adenoma | 77 | 1 | ERR688629 |
| 530890 | carcinoma | 77 | 1 | ERR688634 |
| 531128 | carcinoma | 52 | 1 | ERR688635 |

| | | | | |
|---|---|---|---|---|
| 531155 | carcinoma | 72 | 2 | ERR688636 |
| 531248 | carcinoma | 79 | 1 | ERR688637 |
| 531274 | carcinoma | 77 | 1 | ERR688638 |
| 531277 | carcinoma | 64 | 1 | ERR688639 |
| 531281 | carcinoma | 72 | 2 | ERR688640 |
| 531333 | carcinoma | 69 | 1 | ERR688641 |
| 531352 | carcinoma | 66 | 1 | ERR688642 |
| 531361 | carcinoma | 72 | 2 | ERR688643 |
| 531382 | carcinoma | 65 | 1 | ERR688644 |
| 531416 | carcinoma | 63 | 1 | ERR688647 |
| 531469 | carcinoma | 45 | 2 | ERR688649 |
| 531766 | carcinoma | 72 | 1 | ERR688650 |
| 532832 | advanced adenoma | 68 | 2 | ERR710431 |
