## Supplementary material for "The virus integrations in gut microbes associated with dysbiosis of microbial community in tumorigenesis of colorectal carcinoma": Table S2 (1)

**Table S2. The phage taxonomy of the genetic integration in carcinoma patients.**

| Species | Accession | Regnum | Group |
|---|---|---|---|
| Aureococcu anophagefferen viru | NC_024697 | Virue | dsDNA viruses, no RNA stage |
| Bacteroide phage B124-14 | NC_016770 | Virue | dsDNA viruses, no RNA stage |
| Bacteroide phage B40-8 | NC_011222 | Virue | dsDNA viruses, no RNA stage |
| Cronobacter phage ENT39118 | NC_019934 | Virue | dsDNA viruses, no RNA stage |
| Cronobacter phage ENT47670 | NC_019927 | Virue | dsDNA viruses, no RNA stage |
| Enterobacterial phage mEp213 | NC_019720 | Virue | dsDNA viruses, no RNA stage |
| Enterobacterial phage mEp234 | NC_019715 | Virue | dsDNA viruses, no RNA stage |
| Enterobacterial phage mEp390 | NC_019721 | Virue | dsDNA viruses, no RNA stage |
| Enterobacteria phage 933W | NC_000924 | Virue | dsDNA viruses, no RNA stage |
| Enterobacteria phage BP-4795 | NC_004813 | Virue | dsDNA viruses, no RNA stage |
| Enterobacteria phage cdtI | NC_009514 | Virue | dsDNA viruses, no RNA stage |
| Enterobacteria phage fiAA91- | NC_022750 | Virue | dsDNA viruses, no RNA stage |
| Enterobacteria phage HK106 | NC_019768 | Virue | dsDNA viruses, no RNA stage |
| Enterobacteria phage HK140 | NC_019710 | Virue | dsDNA viruses, no RNA stage |
| Enterobacteria phage HK225 | NC_019717 | Virue | dsDNA viruses, no RNA stage |
| Enterobacteria phage HK446 | NC_019714 | Virue | dsDNA viruses, no RNA stage |
| Enterobacteria phage HK542 | NC_019769 | Virue | dsDNA viruses, no RNA stage |
| Enterobacteria phage HK544 | NC_019767 | Virue | dsDNA viruses, no RNA stage |
| Enterobacteria phage HK629 | NC_019711 | Virue | dsDNA viruses, no RNA stage |
| Enterobacteria phage HK630 | NC_019723 | Virue | dsDNA viruses, no RNA stage |
| Enterobacteria phage HK633 | NC_019719 | Virue | dsDNA viruses, no RNA stage |
| Enterobacteria phage IME10 | NC_019501 | Virue | dsDNA viruses, no RNA stage |
| Enterobacteria phage mEp043 c-1 | NC_019706 | Virue | dsDNA viruses, no RNA stage |
| Enterobacteria phage mEp235 | NC_019708 | Virue | dsDNA viruses, no RNA stage |
| Enterobacteria phage mEp237 | NC_019704 | Virue | dsDNA viruses, no RNA stage |
| Enterobacteria phage mEp460 | NC_019716 | Virue | dsDNA viruses, no RNA stage |
| Enterobacteria phage mEpX1 | NC_019709 | Virue | dsDNA viruses, no RNA stage |
| Enterobacteria phage mEpX2 | NC_019705 | Virue | dsDNA viruses, no RNA stage |
| Enterobacteria phage P4 | NC_001609 | Virue | dsDNA viruses, no RNA stage |
| Enterobacteria phage P88 | NC_026014 | Virue | dsDNA viruses, no RNA stage |
| Enterobacteria phage Phi1 | NC_009821 | Virue | dsDNA viruses, no RNA stage |
| Enterobacteria phage SfV | NC_003444 | Virue | dsDNA viruses, no RNA stage |
| Enterobacteria phage ST104 | NC_005841 | Virue | dsDNA viruses, no RNA stage |
| Enterobacteria phage VT2-Sakai | NC_000902 | Virue | dsDNA viruses, no RNA stage |
| Enterobacteria phage YYZ-2008 | NC_011356 | Virue | unclaified bacterial virue. |
| Enterococcu phage phiEf11 | NC_013696 | Virue | dsDNA viruses, no RNA stage |
| Enterococcu phage phiFL1A | NC_013646 | Virue | dsDNA viruses, no RNA stage |
| Enterococcu phage phiFL2A | NC_013643 | Virue | dsDNA viruses, no RNA stage |
| Enterococcu phage phiFL3A | NC_013648 | Virue | dsDNA viruses, no RNA stage |
| Enterococcu phage phiFL4A | NC_013644 | Virue | dsDNA viruses, no RNA stage |
| Enterococcu phage vB_EfaS_IME197 | NC_028671 | Virue | dsDNA viruses, no RNA stage |
| Echerichia phage D108 | NC_013594 | Virue | dsDNA viruses, no RNA stage |
| Echerichia phage HK639 | NC_016158 | Virue | dsDNA viruses, no RNA stage |
| Echerichia phage HK75 | NC_016160 | Virue | dsDNA viruses, no RNA stage |
| Echerichia phage Min27 | NC_010237 | Virue | dsDNA viruses, no RNA stage |
| Echerichia phage phiV10 | NC_007804 | Virue | dsDNA viruses, no RNA stage |
| Echerichia phage TL-2011b | NC_019445 | Virue | dsDNA viruses, no RNA stage |
| Echerichia phage wV8 | NC_012749 | Virue | dsDNA viruses, no RNA stage |

| | | | |
|---|---|---|---|
| Echerichia Stx1 converting phage | NC_004913 | Virue | dsDNA viruses, no RNA stage |
| Echerichia viru HK022 | NC_002166 | Virue | dsDNA viruses, no RNA stage |
| Echerichia viru HK97 | NC_002167 | Virue | dsDNA viruses, no RNA stage |
| Echerichia viru Lambda | NC_001416 | Virue | dsDNA viruses, no RNA stage |
| Echerichia viru Mu | NC_000929 | Virue | dsDNA viruses, no RNA stage |
| Echerichia viru P1 | NC_005856 | Virue | dsDNA viruses, no RNA stage |
| Echerichia viru P2 | NC_001895 | Virue | dsDNA viruses, no RNA stage |
| Klebiella phage phiKO2 | NC_005857 | Virue | dsDNA viruses, no RNA stage |
| Lactobacillu phage A2 | NC_004112 | Virue | dsDNA viruses, no RNA stage |
| Lactobacillu phage J-1 | NC_022756 | Virue | dsDNA viruses, no RNA stage |
| Lactobacillu phage Lc-Nu | NC_007501 | Virue | dsDNA viruses, no RNA stage |
| Lactobacillu phage phiAT3 | NC_005893 | Virue | dsDNA viruses, no RNA stage |
| Lactobacillu phage phi jlb1 | NC_024206 | Virue | dsDNA viruses, no RNA stage |
| Lactobacillu prophage Lj965 | NC_005355 | Virue | dsDNA viruses, no RNA stage |
| Lactococcu phage 340 | NC_021853 | Virue | dsDNA viruses, no RNA stage |
| Lactococcu phage bIL286 | NC_002667 | Virue | dsDNA viruses, no RNA stage |
| Lactococcu phage BM13 | NC_021861 | Virue | dsDNA viruses, no RNA stage |
| Lactococcu phage jm2 | NC_021860 | Virue | dsDNA viruses, no RNA stage |
| Lactococcu phage jm3 | NC_021854 | Virue | dsDNA viruses, no RNA stage |
| Lactococcu phage P680 | NC_021852 | Virue | dsDNA viruses, no RNA stage |
| Lactococcu phage phi7 | NC_021855 | Virue | dsDNA viruses, no RNA stage |
| Lactococcu phage Tuc2009 | NC_002703 | Virue | dsDNA viruses, no RNA stage |
| Lactococcu viru 712 | NC_008370 | Virue | dsDNA viruses, no RNA stage |
| Lactococcu viru Bibb29 | NC_011046 | Virue | dsDNA viruses, no RNA stage |
| Lactococcu viru bIL170 | NC_001909 | Virue | dsDNA viruses, no RNA stage |
| Lactococcu viru jj50 | NC_008371 | Virue | dsDNA viruses, no RNA stage |
| Lactococcu viru P008 | NC_008363 | Virue | dsDNA viruses, no RNA stage |
| Lactococcu viru k1 | NC_001835 | Virue | dsDNA viruses, no RNA stage |
| Salmonella phage epilon34 | NC_011976 | Virue | dsDNA viruses, no RNA stage |
| Salmonella phage FelixO1 | NC_005282 | Virue | dsDNA viruses, no RNA stage |
| Salmonella phage Fel-2 | NC_010463 | Virue | dsDNA viruses, no RNA stage |
| Salmonella phage g341c | NC_013059 | Virue | dsDNA viruses, no RNA stage |
| Salmonella phage HK620 | NC_002730 | Virue | dsDNA viruses, no RNA stage |
| Salmonella phage RE-2010 | NC_019488 | Virue | dsDNA viruses, no RNA stage |
| Salmonella phage SE1 | NC_011802 | Virue | dsDNA viruses, no RNA stage |
| Salmonella phage SPN3UB | NC_019545 | Virue | dsDNA viruses, no RNA stage |
| Salmonella phage SPN9CC | NC_017985 | Virue | dsDNA viruses, no RNA stage |
| Salmonella phage ST160 | NC_014900 | Virue | dsDNA viruses, no RNA stage |
| Salmonella phage ST64T | NC_004348 | Virue | dsDNA viruses, no RNA stage |
| Salmonella phage vB_SemP_Emek | NC_018275 | Virue | dsDNA viruses, no RNA stage |
| Salmonella phage vB_SoS_Olo | NC_018279 | Virue | dsDNA viruses, no RNA stage |
| Salmonella phage Vi II-E1 | NC_010495 | Virue | dsDNA viruses, no RNA stage |
| Salmonella viru P22 | NC_002371 | Virue | dsDNA viruses, no RNA stage |
| Shigella phage Sf6 | NC_005344 | Virue | dsDNA viruses, no RNA stage |
| Shigella phage SfII | NC_021857 | Virue | dsDNA viruses, no RNA stage |
| Shigella phage SfIV | NC_022749 | Virue | dsDNA viruses, no RNA stage |
| Streptococcu phage 5093 | NC_012753 | Virue | dsDNA viruses, no RNA stage |
| Streptococcu phage M102 | NC_012884 | Virue | dsDNA viruses, no RNA stage |
| Streptococcu phage TP-J34 | NC_020197 | Virue | dsDNA viruses, no RNA stage |
| Stx2-converting phage 1717 | NC_011357 | Virue | dsDNA viruses, no RNA stage |
| Stx2-converting phage 86 | NC_008464 | Virue | dsDNA viruses, no RNA stage |

| | | | |
|---|---|---|---|
| unCultured crAphage | NC_024711 | Virue | unclaified bacterial virue |
| White pot yndrome viru | NC_003225 | Virue | dsDNA viruses, no RNA stage |
| Yerinia phage L-413C | NC_004745 | Virue | dsDNA viruses, no RNA stage |

| Ordo | Familia |
|---|---|
| Phycodnaviridae | unclaifiedPhycodnaviridae. |
| Caudovirale | Siphoviridae. |
| Caudovirale | Siphoviridae. |
| Caudovirale | Siphoviridae. |
| Caudovirale | Myoviridae. |
| Caudovirale | Siphoviridae |
| Caudovirale | Siphoviridae |
| Caudovirale | Siphoviridae |
| Caudovirale | Podoviridae |
| Caudovirale | Siphoviridae |
| Caudovirale | Siphoviridae |
| Caudovirale | Myoviridae |
| Caudovirale | Siphoviridae. |
| Caudovirale | Siphoviridae |
| Caudovirale | Siphoviridae |
| Caudovirale | Siphoviridae |
| Caudovirale | Siphoviridae. |
| Caudovirale | Siphoviridae. |
| Caudovirale | Siphoviridae |
| Caudovirale | Siphoviridae |
| Caudovirale | Siphoviridae |
| Caudovirale | Podoviridae. |
| Caudovirale | Siphoviridae |
| Caudovirale | Siphoviridae |
| Caudovirale | Siphoviridae |
| Caudovirale | Siphoviridae |
| Caudovirale | Siphoviridae |
| Caudovirale | Siphoviridae |
| Caudovirale | Myoviridae |
| Caudovirale | Myoviridae. |
| Caudovirale | Myoviridae |
| Caudovirale | Myoviridae. |
| Caudovirale | Podoviridae |
| Caudovirale | Podoviridae. |
| Caudovirale | Siphoviridae. |
| Caudovirale | Siphoviridae |
| Caudovirale | Siphoviridae |
| Caudovirale | Siphoviridae |
| Caudovirale | Siphoviridae. |
| Caudovirale | Siphoviridae. |
| Caudovirale | Myoviridae |
| Caudovirale | Siphoviridae |
| Caudovirale | Siphoviridae |
| Caudovirale | Podoviridae |
| Caudovirale | Podoviridae. |
| Caudovirale | Podoviridae |
| Caudovirale | Myoviridae |

| | |
|---|---|
| Caudovirale | Podoviridae. |
| Caudovirale | Siphoviridae |
| Caudovirale | Siphoviridae |
| Caudovirale | Siphoviridae |
| Caudovirale | Myoviridae |
| Caudovirale | Myoviridae |
| Caudovirale | Myoviridae |
| Caudovirale | Siphoviridae. |
| Caudovirale | Siphoviridae. |
| Caudovirale | Siphoviridae. |
| Caudovirale | Siphoviridae. |
| Caudovirale | Siphoviridae. |
| Caudovirale | Myoviridae. |
| Caudovirale | Siphoviridae. |
| Caudovirale | Siphoviridae. |
| Caudovirale | Siphoviridae. |
| Caudovirale | Siphoviridae. |
| Caudovirale | Siphoviridae. |
| Caudovirale | Siphoviridae. |
| Caudovirale | Siphoviridae. |
| Caudovirale | Siphoviridae. |
| Caudovirale | Siphoviridae. |
| Caudovirale | Siphoviridae |
| Caudovirale | Siphoviridae |
| Caudovirale | Siphoviridae |
| Caudovirale | Siphoviridae |
| Caudovirale | Siphoviridae |
| Caudovirale | Siphoviridae |
| Caudovirale | Podoviridae |
| Caudovirale | Myoviridae |
| Caudovirale | Myoviridae |
| Caudovirale | Podoviridae |
| Caudovirale | Podoviridae |
| Caudovirale | Myoviridae. |
| Caudovirale | Podoviridae |
| Caudovirale | Siphoviridae. |
| Caudovirale | Podoviridae. |
| Caudovirale | Podoviridae |
| Caudovirale | Podoviridae |
| Caudovirale | Podoviridae |
| Caudovirale | Siphoviridae. |
| Caudovirale | Siphoviridae. |
| Caudovirale | Podoviridae |
| Caudovirale | Podoviridae |
| Caudovirale | Myoviridae. |
| Caudovirale | Myoviridae. |
| Caudovirale | Siphoviridae. |
| Caudovirale | Siphoviridae. |
| Caudovirale | Siphoviridae. |
| Caudovirale | Siphoviridae |
| Caudovirale | Podoviridae. |

| | |
|---|---|
| Nimaviridae | Whipoviru. |
| Caudovirale | Myoviridae |

**Table S2. The phage taxonomy of the genetic integration in advance**

| Species | Accession | Regnum |
|---|---|---|
| Bacteroides phage B124-14 | NC_016770 | Viruses |
| Caviid betaherpesvirus 2 | NC_020231 | Viruses |
| Cronobacter phage ENT39118 | NC_019934 | Viruses |
| Cronobacter phage ENT47670 | NC_019927 | Viruses |
| Enterobacterial phage mEp213 | NC_019720 | Viruses |
| Enterobacteria phage 933W | NC_000924 | Viruses |
| Enterobacteria phage BP-4795 | NC_004813 | Viruses |
| Enterobacteria phage cdtI | NC_009514 | Viruses |
| Enterobacteria phage fiAA91-ss | NC_022750 | Viruses |
| Enterobacteria phage HK106 | NC_019768 | Viruses |
| Enterobacteria phage HK140 | NC_019710 | Viruses |
| Enterobacteria phage HK446 | NC_019714 | Viruses |
| Enterobacteria phage HK629 | NC_019711 | Viruses |
| Enterobacteria phage HK630 | NC_019723 | Viruses |
| Enterobacteria phage HK633 | NC_019719 | Viruses |
| Enterobacteria phage IME10 | NC_019501 | Viruses |
| Enterobacteria phage mEp043 c-1 | NC_019706 | Viruses |
| Enterobacteria phage mEp237 | NC_019704 | Viruses |
| Enterobacteria phage P4 | NC_001609 | Viruses |
| Enterobacteria phage P88 | NC_026014 | Viruses |
| Enterobacteria phage SfV | NC_003444 | Viruses |
| Enterobacteria phage ST104 | NC_005841 | Viruses |
| Enterobacteria phage VT2-Sakai | NC_000902 | Viruses |
| Enterobacteria phage YYZ-2008 | NC_011356 | Viruses |
| Enterococcus phage phiEf11 | NC_013696 | Viruses |
| Enterococcus phage phiFL1A | NC_013646 | Viruses |
| Enterococcus phage phiFL2A | NC_013643 | Viruses |
| Enterococcus phage phiFL3A | NC_013648 | Viruses |
| Enterococcus phage vB_EfaS_IME197 | NC_028671 | Viruses |
| Escherichia phage D108 | NC_013594 | Viruses |
| Escherichia phage HK639 | NC_016158 | Viruses |
| Escherichia phage Min27 | NC_010237 | Viruses |
| Escherichia phage P13374 | NC_018846 | Viruses |
| Escherichia phage phiV10 | NC_007804 | Viruses |
| Escherichia phage Stx2 II | NC_004914 | Viruses |
| Escherichia phage TL-2011b | NC_019445 | Viruses |
| Escherichia phage TL-2011c | NC_019442 | Viruses |
| Escherichia phage wV8 | NC_012749 | Viruses |
| Escherichia Stx1 converting phage | NC_004913 | Viruses |
| Escherichia Stx1 converting phage | NC_004913 | Viruses |
| Escherichia virus 186 | NC_001317 | Viruses |
| Escherichia virus HK022 | NC_002166 | Viruses |
| Escherichia virus Lambda | NC_001416 | Viruses |
| Escherichia virus Mu | NC_000929 | Viruses |
| Escherichia virus P1 | NC_005856 | Viruses |
| Escherichia virus P2 | NC_001895 | Viruses |
| Klebsiella phage phiKO2 | NC_005857 | Viruses |
| Lactobacillus phage A2 | NC_004112 | Viruses |

| | | |
|---|---|---|
| Lactobacillus phage J-1 | NC_022756 | Viruses |
| Lactobacillus phage PL-1 | NC_022757 | Viruses |
| Lactococcus phage bIL285 | NC_002666 | Viruses |
| Lactococcus phage bIL286 | NC_002667 | Viruses |
| Lactococcus phage bIL309 | NC_002668 | Viruses |
| Lactococcus phage bIL310 | NC_002669 | Viruses |
| Lactococcus phage bIL312 | NC_002671 | Viruses |
| Lactococcus phage BK5-T | NC_002796 | Viruses |
| Lactococcus phage BM13 | NC_021861 | Viruses |
| Lactococcus phage jm2 | NC_021860 | Viruses |
| Lactococcus phage P335 sensu lato | NC_004746 | Viruses |
| Lactococcus phage phiL47 | NC_023574 | Viruses |
| Lactococcus phage phiLC3 | NC_005822 | Viruses |
| Lactococcus phage r1t | NC_004302 | Viruses |
| Lactococcus phage TP901-1 | NC_002747 | Viruses |
| Lactococcus phage Tuc2009 | NC_002703 | Viruses |
| Lactococcus phage ul36 | NC_004066 | Viruses |
| Lactococcus virus P008 | NC_008363 | Viruses |
| Salmonella phage epsilon15 | NC_004775 | Viruses |
| Salmonella phage epsilon34 | NC_011976 | Viruses |
| Salmonella phage HK620 | NC_002730 | Viruses |
| Salmonella phage SSU5 | NC_018843 | Viruses |
| Salmonella phage ST160 | NC_014900 | Viruses |
| Salmonella virus P22 | NC_002371 | Viruses |
| Salmonella virus PsP3 | NC_005340 | Viruses |
| Shigella phage Sf6 | NC_005344 | Viruses |
| Shigella phage SfII | NC_021857 | Viruses |
| Shigella phage SfIV | NC_022749 | Viruses |
| Streptococcus phage IC1 | NC_024370 | Viruses |
| Streptococcus phage Sfi21 | NC_000872 | Viruses |
| Streptococcus phage TP-J34 | NC_020197 | Viruses |
| Stx2-converting phage 1717 | NC_011357 | Viruses |
| Stx2-converting phage 86 | NC_008464 | Viruses |
| Vibrio phage pYD38-A | NC_021534 | Viruses |
| Yersinia phage L-413C | NC_004745 | Viruses |

**d_adenoma patients.**

| Group | Ordo | Familia |
|---|---|---|
| dsDNA viruses, no RNA stage | Caudovirales | Siphoviridae. |
| dsDNA viruses, no RNA stage | Herpesvirales | Herpesviridae |
| dsDNA viruses, no RNA stage | Caudovirales | Siphoviridae. |
| dsDNA viruses, no RNA stage | Caudovirales | Myoviridae. |
| dsDNA viruses, no RNA stage | Caudovirales | Siphoviridae |
| dsDNA viruses, no RNA stage | Caudovirales | Podoviridae |
| dsDNA viruses, no RNA stage | Caudovirales | Siphoviridae |
| dsDNA viruses, no RNA stage | Caudovirales | Siphoviridae |
| dsDNA viruses, no RNA stage | Caudovirales | Myoviridae |
| dsDNA viruses, no RNA stage | Caudovirales | Siphoviridae. |
| dsDNA viruses, no RNA stage | Caudovirales | Siphoviridae |
| dsDNA viruses, no RNA stage | Caudovirales | Siphoviridae |
| dsDNA viruses, no RNA stage | Caudovirales | Siphoviridae |
| dsDNA viruses, no RNA stage | Caudovirales | Siphoviridae |
| dsDNA viruses, no RNA stage | Caudovirales | Siphoviridae |
| dsDNA viruses, no RNA stage | Caudovirales | Podoviridae. |
| dsDNA viruses, no RNA stage | Caudovirales | Siphoviridae |
| dsDNA viruses, no RNA stage | Caudovirales | Siphoviridae |
| dsDNA viruses, no RNA stage | Caudovirales | Myoviridae |
| dsDNA viruses, no RNA stage | Caudovirales | Myoviridae. |
| dsDNA viruses, no RNA stage | Caudovirales | Myoviridae. |
| dsDNA viruses, no RNA stage | Caudovirales | Podoviridae |
| dsDNA viruses, no RNA stage | Caudovirales | Podoviridae. |
| unclassified bacterial viruses. | | |
| dsDNA viruses, no RNA stage | Caudovirales | Siphoviridae. |
| dsDNA viruses, no RNA stage | Caudovirales | Siphoviridae |
| dsDNA viruses, no RNA stage | Caudovirales | Siphoviridae |
| dsDNA viruses, no RNA stage | Caudovirales | Siphoviridae |
| dsDNA viruses, no RNA stage | Caudovirales | Siphoviridae. |
| dsDNA viruses, no RNA stage | Caudovirales | Myoviridae |
| dsDNA viruses, no RNA stage | Caudovirales | Siphoviridae |
| dsDNA viruses, no RNA stage | Caudovirales | Podoviridae |
| dsDNA viruses, no RNA stage | Caudovirales | Podoviridae. |
| dsDNA viruses, no RNA stage | Caudovirales | Podoviridae. |
| dsDNA viruses, no RNA stage | Caudovirales | Podoviridae |
| dsDNA viruses, no RNA stage | Caudovirales | Podoviridae |
| dsDNA viruses, no RNA stage | Caudovirales | Podoviridae |
| dsDNA viruses, no RNA stage | Caudovirales | Myoviridae |
| dsDNA viruses, no RNA stage | Caudovirales | Podoviridae. |
| dsDNA viruses, no RNA stage | Caudovirales | Podoviridae. |
| dsDNA viruses, no RNA stage | Caudovirales | Myoviridae |
| dsDNA viruses, no RNA stage | Caudovirales | Siphoviridae |
| dsDNA viruses, no RNA stage | Caudovirales | Siphoviridae |
| dsDNA viruses, no RNA stage | Caudovirales | Myoviridae |
| dsDNA viruses, no RNA stage | Caudovirales | Myoviridae |
| dsDNA viruses, no RNA stage | Caudovirales | Myoviridae |
| dsDNA viruses, no RNA stage | Caudovirales | Siphoviridae. |
| dsDNA viruses, no RNA stage | Caudovirales | Siphoviridae. |

| | | |
|---|---|---|
| dsDNA viruses, no RNA stage | Caudovirales | Siphoviridae. |
| dsDNA viruses, no RNA stage | Caudovirales | Siphoviridae. |
| dsDNA viruses, no RNA stage | Caudovirales | Siphoviridae. |
| dsDNA viruses, no RNA stage | Caudovirales | Siphoviridae. |
| dsDNA viruses, no RNA stage | Caudovirales | Siphoviridae. |
| dsDNA viruses, no RNA stage | Caudovirales | Siphoviridae. |
| dsDNA viruses, no RNA stage | Caudovirales | Siphoviridae. |
| dsDNA viruses, no RNA stage | Caudovirales | Siphoviridae. |
| dsDNA viruses, no RNA stage | Caudovirales | Siphoviridae. |
| dsDNA viruses, no RNA stage | Caudovirales | Siphoviridae. |
| dsDNA viruses, no RNA stage | Caudovirales | Siphoviridae. |
| dsDNA viruses, no RNA stage | Caudovirales | Siphoviridae. |
| dsDNA viruses, no RNA stage | Caudovirales | Siphoviridae. |
| dsDNA viruses, no RNA stage | Caudovirales | Siphoviridae. |
| dsDNA viruses, no RNA stage | Caudovirales | Siphoviridae. |
| dsDNA viruses, no RNA stage | Caudovirales | Siphoviridae. |
| dsDNA viruses, no RNA stage | Caudovirales | Siphoviridae. |
| dsDNA viruses, no RNA stage | Caudovirales | Siphoviridae |
| dsDNA viruses, no RNA stage | Caudovirales | Podoviridae. |
| dsDNA viruses, no RNA stage | Caudovirales | Podoviridae |
| dsDNA viruses, no RNA stage | Caudovirales | Podoviridae |
| dsDNA viruses, no RNA stage | Caudovirales | Siphoviridae. |
| dsDNA viruses, no RNA stage | Caudovirales | Podoviridae |
| dsDNA viruses, no RNA stage | Caudovirales | Podoviridae |
| dsDNA viruses, no RNA stage | Caudovirales | Myoviridae |
| dsDNA viruses, no RNA stage | Caudovirales | Podoviridae |
| dsDNA viruses, no RNA stage | Caudovirales | Myoviridae. |
| dsDNA viruses, no RNA stage | Caudovirales | Myoviridae. |
| unclassified bacterial viruses. | | |
| dsDNA viruses, no RNA stage | Caudovirales | Siphoviridae |
| dsDNA viruses, no RNA stage | Caudovirales | Siphoviridae. |
| dsDNA viruses, no RNA stage | Caudovirales | Siphoviridae |
| dsDNA viruses, no RNA stage | Caudovirales | Podoviridae. |
| dsDNA viruses, no RNA stage | unclassified dsDNA phages. | |
| dsDNA viruses, no RNA stage | Caudovirales | Myoviridae |

**Table S2. The phage taxonomy of the genetic integration in healthy controls.**

| Species | Accession | Regnum | Group |
|---|---|---|---|
| Bacteroides phage B124-14 | NC_016770 | Viruses | dsDNA viruses, no RNA stage |
| Enterobacteria phage 933W | NC_000924 | Viruses | dsDNA viruses, no RNA stage |
| Enterobacteria phage BP-4795 | NC_004813 | Viruses | dsDNA viruses, no RNA stage |
| Enterobacteria phage fiAA91-ss | NC_022750 | Viruses | dsDNA viruses, no RNA stage |
| Enterobacteria phage HK106 | NC_019768 | Viruses | dsDNA viruses, no RNA stage |
| Enterobacteria phage HK140 | NC_019710 | Viruses | dsDNA viruses, no RNA stage |
| Enterobacteria phage HK225 | NC_019717 | Viruses | dsDNA viruses, no RNA stage |
| Enterobacteria phage HK446 | NC_019714 | Viruses | dsDNA viruses, no RNA stage |
| Enterobacteria phage HK542 | NC_019769 | Viruses | dsDNA viruses, no RNA stage |
| Enterobacteria phage HK544 | NC_019767 | Viruses | dsDNA viruses, no RNA stage |
| Enterobacteria phage HK629 | NC_019711 | Viruses | dsDNA viruses, no RNA stage |
| Enterobacteria phage HK633 | NC_019719 | Viruses | dsDNA viruses, no RNA stage |
| Enterobacteria phage IME10 | NC_019501 | Viruses | dsDNA viruses, no RNA stage |
| Enterobacteria phage mEp043 c-1 | NC_019706 | Viruses | dsDNA viruses, no RNA stage |
| Enterobacteria phage mEp237 | NC_019704 | Viruses | dsDNA viruses, no RNA stage |
| Enterobacteria phage mEpX1 | NC_019709 | Viruses | dsDNA viruses, no RNA stage |
| Enterobacteria phage mEpX2 | NC_019705 | Viruses | dsDNA viruses, no RNA stage |
| Enterobacteria phage P88 | NC_026014 | Viruses | dsDNA viruses, no RNA stage |
| Enterobacteria phage YYZ-2008 | NC_011356 | Viruses | u,NClassified bacterial viruses. |
| Enterococcus phage phiFL3A | NC_013648 | Viruses | dsDNA viruses, no RNA stage |
| Escherichia phage HK639 | NC_016158 | Viruses | dsDNA viruses, no RNA stage |
| Escherichia phage phiV10 | NC_007804 | Viruses | dsDNA viruses, no RNA stage |
| Escherichia phage TL-2011b | NC_019445 | Viruses | dsDNA viruses, no RNA stage |
| Escherichia Stx1 converting phage | NC_004913 | Viruses | dsDNA viruses, no RNA stage |
| Escherichia virus 186 | NC_001317 | Viruses | dsDNA viruses, no RNA stage |
| Escherichia virus HK97 | NC_002167 | Viruses | dsDNA viruses, no RNA stage |
| Escherichia virus Lambda | NC_001416 | Viruses | dsDNA viruses, no RNA stage |
| Escherichia virus Mu | NC_000929 | Viruses | dsDNA viruses, no RNA stage |
| Escherichia virus P1 | NC_005856 | Viruses | dsDNA viruses, no RNA stage |
| Lactobacillus phage A2 | NC_004112 | Viruses | dsDNA viruses, no RNA stage |
| Lactobacillus phage J-1 | NC_022756 | Viruses | dsDNA viruses, no RNA stage |
| Lactobacillus phage KC5a | NC_007924 | Viruses | dsDNA viruses, no RNA stage |
| Lactobacillus phage Lc-Nu | NC_007501 | Viruses | dsDNA viruses, no RNA stage |
| Lactobacillus phage phiAT3 | NC_005893 | Viruses | dsDNA viruses, no RNA stage |
| Lactobacillus phage PL-1 | NC_022757 | Viruses | dsDNA viruses, no RNA stage |
| Lactococcus phage bIL286 | NC_002667 | Viruses | dsDNA viruses, no RNA stage |
| Lactococcus phage BM13 | NC_021861 | Viruses | dsDNA viruses, no RNA stage |
| Lactococcus phage jm2 | NC_021860 | Viruses | dsDNA viruses, no RNA stage |
| Lactococcus phage jm3 | NC_021854 | Viruses | dsDNA viruses, no RNA stage |
| Lactococcus phage r1t | NC_004302 | Viruses | dsDNA viruses, no RNA stage |
| Lactococcus phage TP901-1 | NC_002747 | Viruses | dsDNA viruses, no RNA stage |
| Lactococcus phage Tuc2009 | NC_002703 | Viruses | dsDNA viruses, no RNA stage |
| Lactococcus phage ul36 | NC_004066 | Viruses | dsDNA viruses, no RNA stage |
| Salmonella phage epsilon34 | NC_011976 | Viruses | dsDNA viruses, no RNA stage |
| Salmonella phage HK620 | NC_002730 | Viruses | dsDNA viruses, no RNA stage |
| Salmonella phage SPN9CC | NC_017985 | Viruses | dsDNA viruses, no RNA stage |
| Salmonella phage SSU5 | NC_018843 | Viruses | dsDNA viruses, no RNA stage |
| Salmonella phage vB_SemP_Emek | NC_018275 | Viruses | dsDNA viruses, no RNA stage |

| | | | |
|---|---|---|---|
| Salmonella phage vB_SosS_Oslo | NC_018279 | Viruses | dsDNA viruses, no RNA stage |
| Salmonella virus P22 | NC_002371 | Viruses | dsDNA viruses, no RNA stage |
| Salmonella virus PsP3 | NC_005340 | Viruses | dsDNA viruses, no RNA stage |
| Streptococcus phage 20617 | NC_023503 | Viruses | u,NClassified bacterial viruses. |
| Streptococcus phage IC1 | NC_024370 | Viruses | u,NClassified bacterial viruses. |
| Streptococcus phage M102 | NC_012884 | Viruses | dsDNA viruses, no RNA stage |
| Streptococcus phage MM1 | NC_003050 | Viruses | dsDNA viruses, no RNA stage |
| Streptococcus phage O1205 | NC_004303 | Viruses | dsDNA viruses, no RNA stage |
| Streptococcus phage Sfi21 | NC_000872 | Viruses | dsDNA viruses, no RNA stage |
| Streptococcus phage TP-J34 | NC_020197 | Viruses | dsDNA viruses, no RNA stage |
| Stx2-converting phage 1717 | NC_011357 | Viruses | dsDNA viruses, no RNA stage |

| Ordo | Familia |
|---|---|
| Caudovirales | Siphoviridae. |
| Caudovirales | Podoviridae |
| Caudovirales | Siphoviridae |
| Caudovirales | Myoviridae |
| Caudovirales | Siphoviridae. |
| Caudovirales | Siphoviridae |
| Caudovirales | Siphoviridae |
| Caudovirales | Siphoviridae |
| Caudovirales | Siphoviridae. |
| Caudovirales | Siphoviridae. |
| Caudovirales | Siphoviridae |
| Caudovirales | Siphoviridae |
| Caudovirales | Podoviridae. |
| Caudovirales | Siphoviridae |
| Caudovirales | Siphoviridae |
| Caudovirales | Siphoviridae |
| Caudovirales | Siphoviridae |
| Caudovirales | Myoviridae. |
| Caudovirales | Siphoviridae |
| Caudovirales | Siphoviridae |
| Caudovirales | Podoviridae. |
| Caudovirales | Podoviridae |
| Caudovirales | Podoviridae. |
| Caudovirales | Myoviridae |
| Caudovirales | Siphoviridae |
| Caudovirales | Siphoviridae |
| Caudovirales | Myoviridae |
| Caudovirales | Myoviridae |
| Caudovirales | Siphoviridae. |
| Caudovirales | Siphoviridae. |
| Caudovirales | Myoviridae. |
| Caudovirales | Siphoviridae. |
| Caudovirales | Siphoviridae. |
| Caudovirales | Siphoviridae. |
| Caudovirales | Siphoviridae. |
| Caudovirales | Siphoviridae. |
| Caudovirales | Siphoviridae. |
| Caudovirales | Siphoviridae. |
| Caudovirales | Siphoviridae. |
| Caudovirales | Siphoviridae. |
| Caudovirales | Siphoviridae. |
| Caudovirales | Siphoviridae. |
| Caudovirales | Podoviridae |
| Caudovirales | Podoviridae |
| Caudovirales | Podoviridae. |
| Caudovirales | Siphoviridae. |
| Caudovirales | Podoviridae |

| | |
|---|---|
| Caudovirales | Siphoviridae. |
| Caudovirales | Podoviridae |
| Caudovirales | Myoviridae |
| Caudovirales | Siphoviridae. |
| Caudovirales | Siphoviridae. |
| Caudovirales | Siphoviridae |
| Caudovirales | Siphoviridae |
| Caudovirales | Siphoviridae. |
| Caudovirales | Siphoviridae |
